# A framework to identify structured behavioral patterns within rodent spatial trajectories

**DOI:** 10.1101/2020.03.02.967489

**Authors:** Francesco Donnarumma, Roberto Prevete, Domenico Maisto, Simone Fuscone, Emily M. Irvine, Matthijs A. A. van der Meer, Caleb Kemere, Giovanni Pezzulo

## Abstract

Animal behavior is highly structured. Yet, structured behavioral patterns – or “statistical ethograms” – are not immediately apparent from the full spatiotemporal data that behavioral scientists usually collect. Here, we introduce a framework to quantitatively characterize rodent behavior during spatial (e.g., maze) navigation, in terms of movement building blocks or *motor primitives*. The hypothesis that we pursue is that rodent behavior is characterized by a small number of motor primitives, which are combined over time to produce open-ended movements. We assume motor primitives to be organized in terms of two sparsity principles: each movement is controlled using a limited subset of motor primitives (sparse superposition) and each primitive is active only for time-limited, time-contiguous portions of movements (sparse activity). We formalize this hypothesis using a sparse dictionary learning method, which we use to extract motor primitives from rodent position and velocity data collected during spatial navigation, and successively to reconstruct past trajectories and predict novel ones. Three main results validate our approach. First, rodent behavioral trajectories are robustly reconstructed from incomplete data, performing better than approaches based on standard dimensionality reduction methods, such as principal component analysis, or single sparsity. Second, the motor primitives extracted during one experimental session generalize and afford the accurate reconstruction of rodent behavior across successive experimental sessions in the same or in modified mazes. Third, in our approach the number of motor primitives associated with each maze correlates with independent measures of maze complexity, hence showing that our formalism is sensitive to essential aspects of task structure. The framework introduced here can be used by behavioral scientists and neuroscientists as an aid for behavioral and neural data analysis. Indeed, the extracted motor primitives enable the quantitative characterization of the complexity and similarity between different mazes and behavioral patterns across multiple trials (i.e., habit formation). We provide example uses of this computational framework, showing how it can be used to identify behavioural effects of maze complexity, analyze stereotyped behavior, classify behavioral choices and predict place and grid cell displacement in novel environments.

## Introduction

During cognitive tasks, such as spatial navigation, animals exhibit highly structured behavior. These behavioral patterns or motifs can indicate important latent dimensions, such as task complexity (e.g., movement variability) or the amount of training that the animal has received (e.g., movement stereotypy). Yet, a standard methodology to extract structured behavioral patterns from the set of data (e.g., position and velocity) that behavioral scientists usually collect during spatial navigation is still lacking.

Here we present a computational framework that characterizes the latent structure – or a “statistical ethogram” – of rodent movement dynamics during spatial navigation, in terms of movement building blocks or motor primitives. The idea that motor control uses a combination of motor primitives is widely accepted in computational neuroscience^1–8^ but its application for the study of rodent spatial navigation is less common. A line of research focusing on rat behavior used a descriptive approach to the study of rodent exploration, which revealed its organization into movement subsystems that can be readily quantified^9–12^. This body of work provides evidence for an organization of behavior in terms of motor primitives, also providing a developmental perspective on their acquisition^13,14^.

### Using motor primitives to study rodent movements during spatial navigation

Motor primitives can be characterized as spatiotemporal patterns that recur across an animal’s behavior and thus permit reconstructing it in a parsimonious way – in the sense that a small number of motor primitives permit reconstructing a large dataset of movements. Figure 1 illustrates schematically the basic elements of the motor primitives approach. Spatial (x,y) and velocity (x,y) data are collected from animal trajectories during spatial navigation (Figure 1a) and used to derive a set (or dictionary) of motor primitives, such as the ones shown in Figure 1b. Later, these motor primitives can be used to “reconstruct” actual animal trajectories (see Figure 1c for an example of reconstruction of spatial components of the trajectories). Figure 1d shows an example of motor primitives superimposed on one of the mazes that we used in our study (see later for explanation).

There are two fundamental (biological and computational) motivations for using a motor primitive approach to study complex behavior. From a biological perspective, the hypothesis that the brain may adapt motor primitives as fundamental units to organize complex (spatial or other) movements in time has received increased attention^15^. In this perspective, complex movements can be composed by combining over time a limited basis of dynamic motor primitives – such that, for each time frame, only one or a few motor primitives are active. Therefore, inferring the animal’s motor primitives during movements can help understanding how the brain organizes complex behavioral patterns.

**Figure 1.**
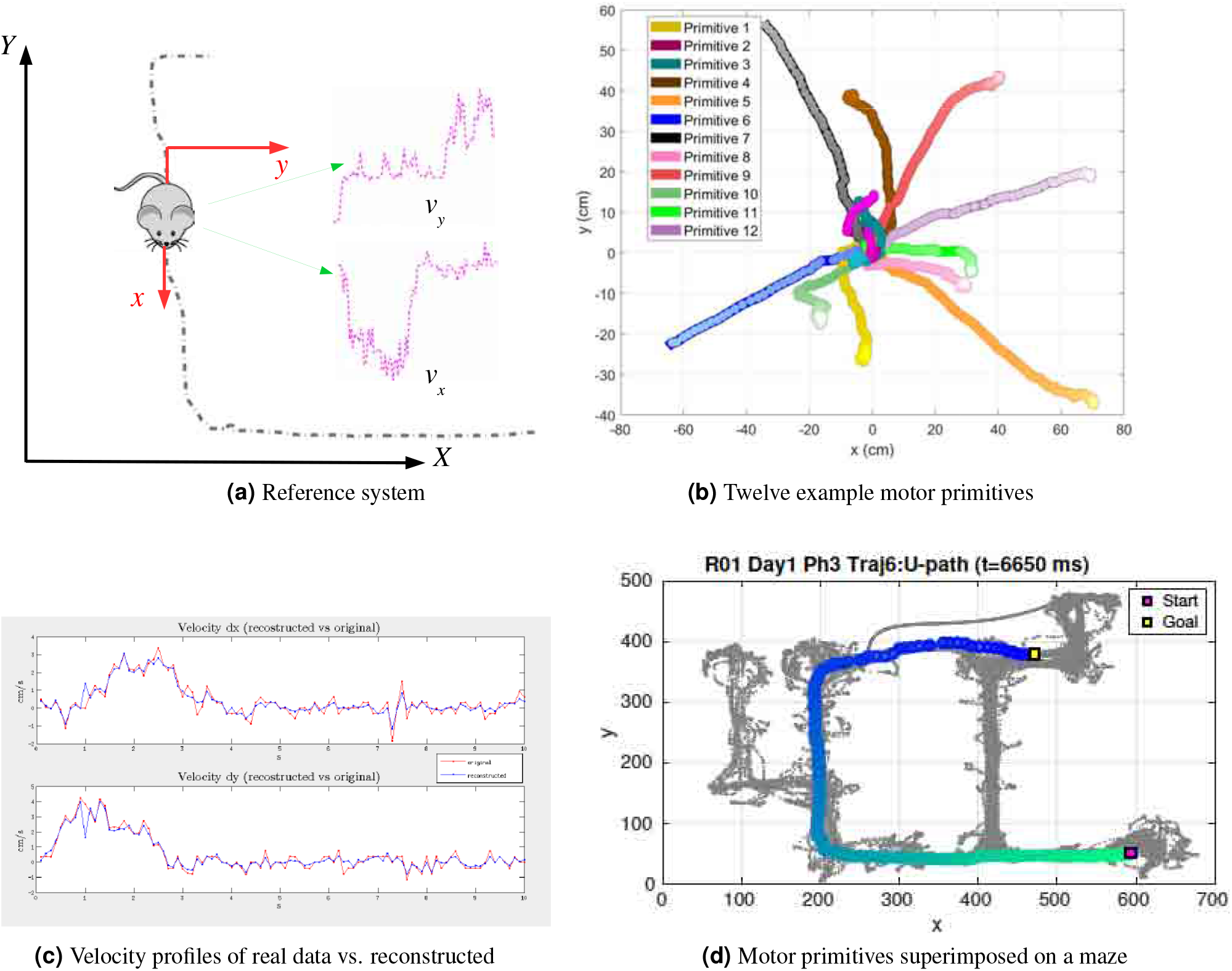
Schematic illustration of the approach. Panel (a): the rodent-centered coordinate system used in input for the dictionary learning model. The dotted black line is the actual animal trajectory. For each time, we extracted from data a 2-D (x, y) spatial position. Note that our reference system is not anchored to the maze. Rather, at each time point, x represents the last direction of the animal. The resulting, animal-centered reference system permits deriving different primitives for different orientations of the animal. Furthermore, we extracted animal velocity in x and y dimensions. Panel (b): Twelve sample motor primitives extracted from navigation data. For illustration purposes we simulated the primitives starting at point (0,0) at time 0. The gradient of lighter colors for circle markers indicates increase in time. Panel (c): example reconstruction of real rodent data using motor primitives (here, only the velocity in x and y axes is reconstructed). Blue is the actual velocity profile over time, red is the reconstructed profile. Panel (d): example superimposition of motor primitives on one of the mazes used in this study. See the main text for explanation.

There is also a second, more practical reason to use the motor primitives approach to study animal movements (even if ultimately the brain uses a different organization). Characterizing animal behavior in terms of its underlying motor primitives can be more practical than using the full spatiotemporal [x(t),y(t)] data and more indicative of underlying regularities (e.g., behavioral stereotypy), as demonstrated by the analysis of behavioral motifs in various animal, species such as Caenorhabditis elegans^16^, fruit fly^17^, mice^18^ and rat^19^. Although previous work has analyzed behaviors such as path stereotypy in terms of path similarity (e.g.^20^), such descriptive approaches cannot generate novel data. More recent unsupervised learning methods^21^ or related approaches^22–26^ are more similar to our approach, but have not yet been applied to structured maze data.

### Double sparsity in the organization of motor primitives and in dictionary learning methods

There are various ways to construct motor primitives at neural and computational level (see Figure 2). The approach pursued here is motivated by a *double sparsity* hypothesis, according to which the organization of motor primitives shows simultaneously two kinds of sparsity, which we call *sparse superposition* and *sparse activity*. *Sparse superposition* implies that at each time step, a movement can be reconstructed by superimposing at most a few primitives – and possibly just one (see Figure 2b). *Sparse activity* entails that the same motor primitive is only active for a time-limited, time-contiguous portion of the movement (see Figure 2c).

**Figure 2.**
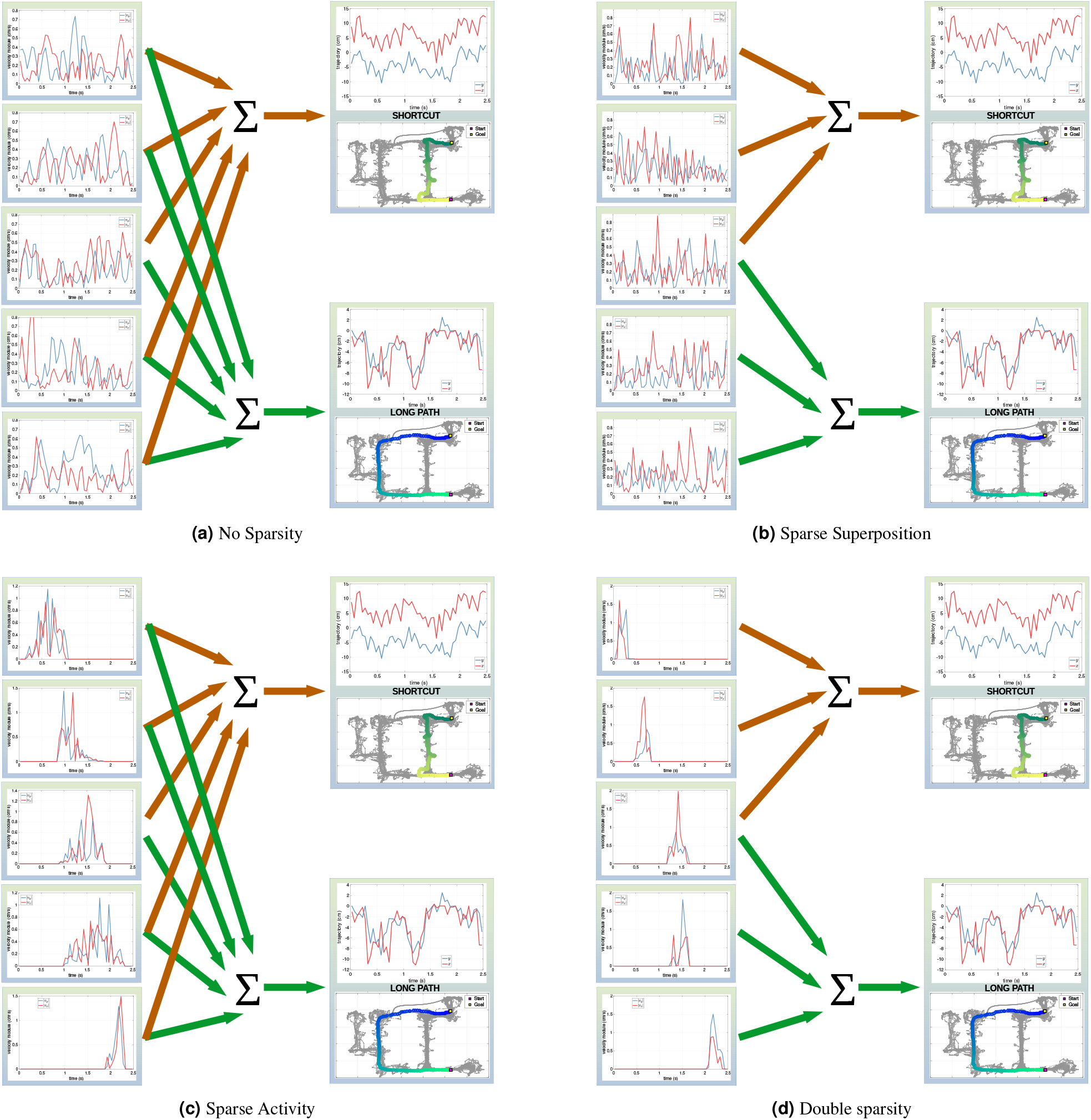
*Four alternative approaches and hypotheses on the organization of motor primitives.* Panel (a) illustrates the hypothesis of no sparsity in motor primitives organization, implemented using Principal Component Analysis (PCA). Panel (b) illustrates the sparse superposition hypothesis, implemented using *ℓ*1-regularization. In this perspective, different actions are represented by a combination of small subsets of primitives, which can overlap. The Figure shows an example with five motor primitives, where the first three are jointly used to represent one motor action, and the last three are jointly used to represent another motor action. Panel (c) illustrates the sparse activity hypothesis, implemented using SSPCA. In this perspective, actions are represented using all available motor primitives, but these are active only in a restricted time interval, rather than for the entire duration of the action. The Figure shows an example with five motor primitives, all of which are used to represent both motor actions, but are time-localized. Panel (d) illustrates the double sparsity hypothesis, which combines the previous two hypotheses and is implemented using SRSSD. See the main text for details.

Figure 2 compares four hypotheses on how motor primitives are organized. The first hypothesis assumes that there is no sparsity in the organization of motor primitives: motor actions are represented as combinations of all the available primitives, for the entire duration of the motor action (Figure 2a). The second hypothesis assumes sparse superposition: given a large enough set of primitives, different actions can be represented by a combination of small subsets of primitives, which can overlap^27,28^ (Figure 2b). The third hypothesis assumes sparse activity: each primitive is active only in a limited time interval, rather than for the entire duration of a motor action^29,30^ (Figure 2c). Finally, the fourth hypothesis combines the second and third hypotheses and assumes dual sparsity: motor actions are represented by combinations of small subsets of primitives, which are only active in a limited time interval (Figure 2d).

The four hypotheses listed above correspond to four computational approaches to identify motor primitives from data. The first approach corresponds directly to Principal Component Analysis (PCA), which does not assume any sparsity. Interestingly, the other three approaches correspond directly to three sparse dictionary learning methods in machine learning^31–33^, which assume different kinds of sparsity: namely, the *ℓ*1-regularization method^34^, which assumes recombination of atoms with *sparse coefficients*, and corresponds to the sparse superposition hypothesis; Structured Sparse Principal Component Analysis (SSPCA)^35^, which assumes *atom structured sparsity* and thus corresponds to the sparse activity hypothesis; and Sparse Representation of Structured Sparse Dictionaries (SRSSD), which assumes double sparsity. In other words, there is a close relation between the concept of sparse motor primitives in biological motor control and methods for sparse dictionary learning in machine learning. In the latter, the equivalent of motor primitives are called *atoms* and they are part of a *dictionary*, which corresponds to the animal’s behavioral repertoire.

In the rest of the article, we introduce a framework for the analysis of rodent movements during spatial navigation that is based on the SRSSD approach – which directly implements a dual sparsity hypothesis. We then compare the SRSSD approach with the three alternative methods, showing that it entails better and more parsimonious reconstruction of data. The possibility to model, predict and reconstruct spatial trajectories in terms of their underlying motor primitives opens the doors to a number of potential applications of the framework, including the possibility to discover sequential structure, repeating patterns and stereotypy in behavior, and predict animal choices – which we discuss and exemplify in the final section.

## Results

The computational framework that we used in this article is illustrated in Figure 3. First, we divided the experimental (rodent trajectory) data in three sets: a training set, a validation set and a test set. We then used the training set to develop several candidate *dictionaries* of motor primitives from data, using *sparse dictionary learning*^31–33^. Furthermore, we evaluated candidate dictionaries on different reconstructions performed on a validation set, in order to select which combination of parameters (*dictionary size*, *coefficient sparsity* and *atom structured sparsity*) afford the best reconstruction of trajectory data. Finally, we used the selected dictionary to reconstruct data of the test set and to simulate real rodent motor data – hence validating the approach and comparing it with alternatives. Below we discuss in more detail all the elements of the framework.

**Figure 3.**
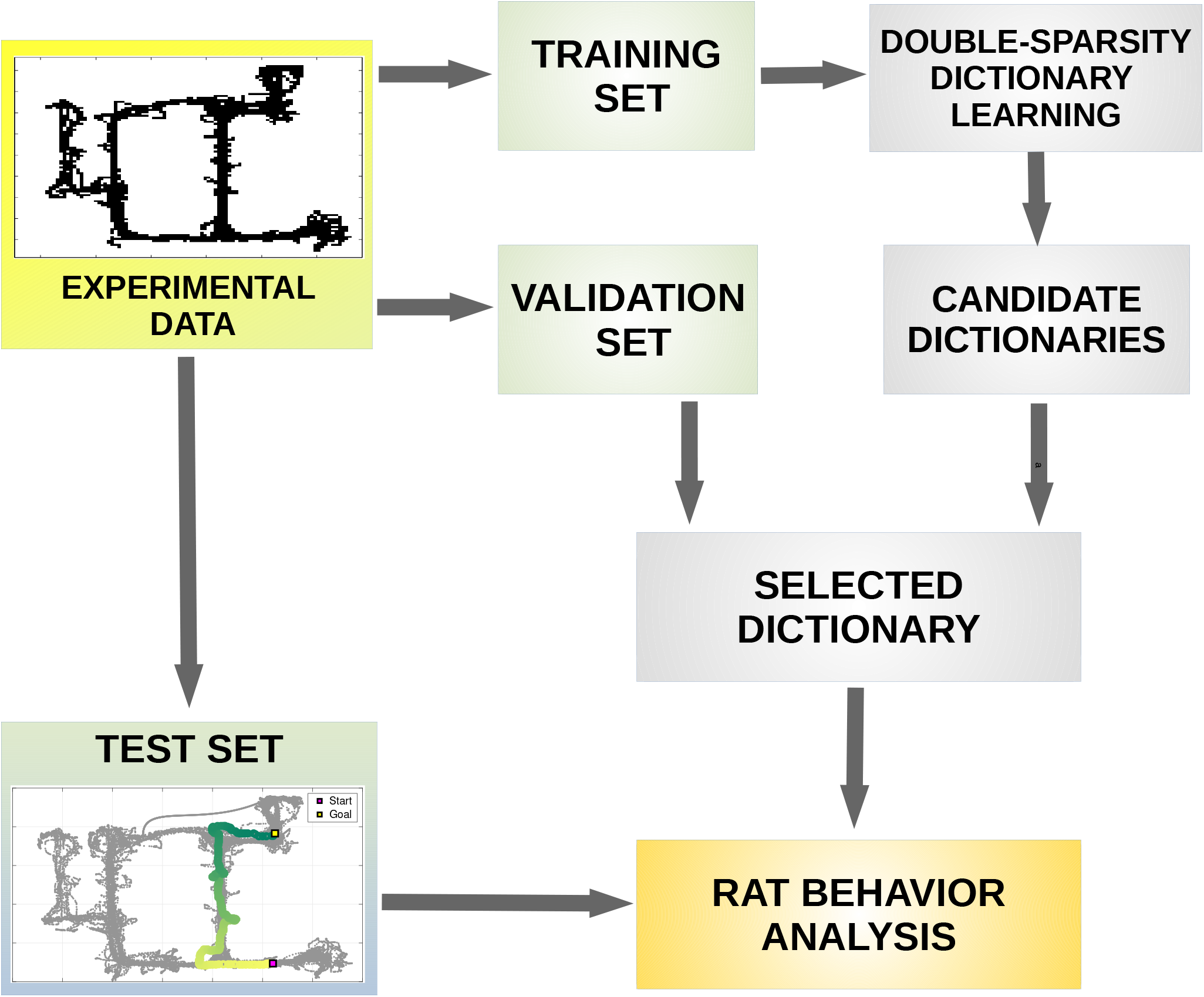
A framework for the analysis of rat behavior. The figure shows the workflow of the framework proposed in our paper. After collecting the experimental data, these data are split into three sets: a training set, a validation set and a test set. The training set is processed by a double-sparse dictionary learning performed with the SRSDD algorithm (see Material and methods Alg. 1), that represents data in a set of atoms - called *dictionary* - which are interpreted as rat motor primitives. Different candidate dictionaries are found during this process with different parameters of atom structured sparsity. Candidate dictionaries are evaluated on different sparse reconstructions performed by means of Alg. 2 on a validation set. At the end of this phase, the selected dictionary can be used on different tests to analyse rat behavior during different motor actions in the maze.

### Dataset

In order to validate our approach, we conducted a series of experiments using a dataset of rodent movements collected in the van der Meer lab (Dartmouth College). This dataset includes data from 3 rats (R01, R02, R03). Each rat experienced a sequence of 8 unique mazes, with novel routes introduced to a familiar environment every session / day. The rats navigated the maze environment between two goal (reward) locations, placed at the ends of a familiar U-shaped track. For each session, maze data was collected in three distinct phases. In the first and second phases, the rats were only able to navigate along the familiar U-shaped track. In the third phase, two novel paths became accessible – one of which was a shortcut between the two reward locations, and the other a dead end. Figure 4 shows the behavioral trajectories from rat R01 for each of the eight mazes in the three phases: phase 1 (green), phase 2 (yellow) and phase 3 (grey). Table 1 shows the overall duration of each of the three phases for rat R01 (across multiple trials, see below) from each of the eight sessions.

**Table 1.** Duration in minutes of the different phases, rat R01

| R01 (min) | Day1 | Day2 | Day3 | Day4 | Day5 | Day6 | Day7 | Day8 |
| --- | --- | --- | --- | --- | --- | --- | --- | --- |
| Phase1 | 10.08 | 10.05 | 8.02 | 8.13 | 9.88 | 8.48 | 8.41 | 8.10 |
| Phase2 | 20.33 | 20.37 | 15.02 | 20.55 | 20.33 | 21.12 | 20.24 | 20.04 |
| Phase3 | 37.01 | 46.10 | 45.05 | 42.68 | 52.21 | 45.82 | 40.43 | 55.04 |

**Figure 4.**
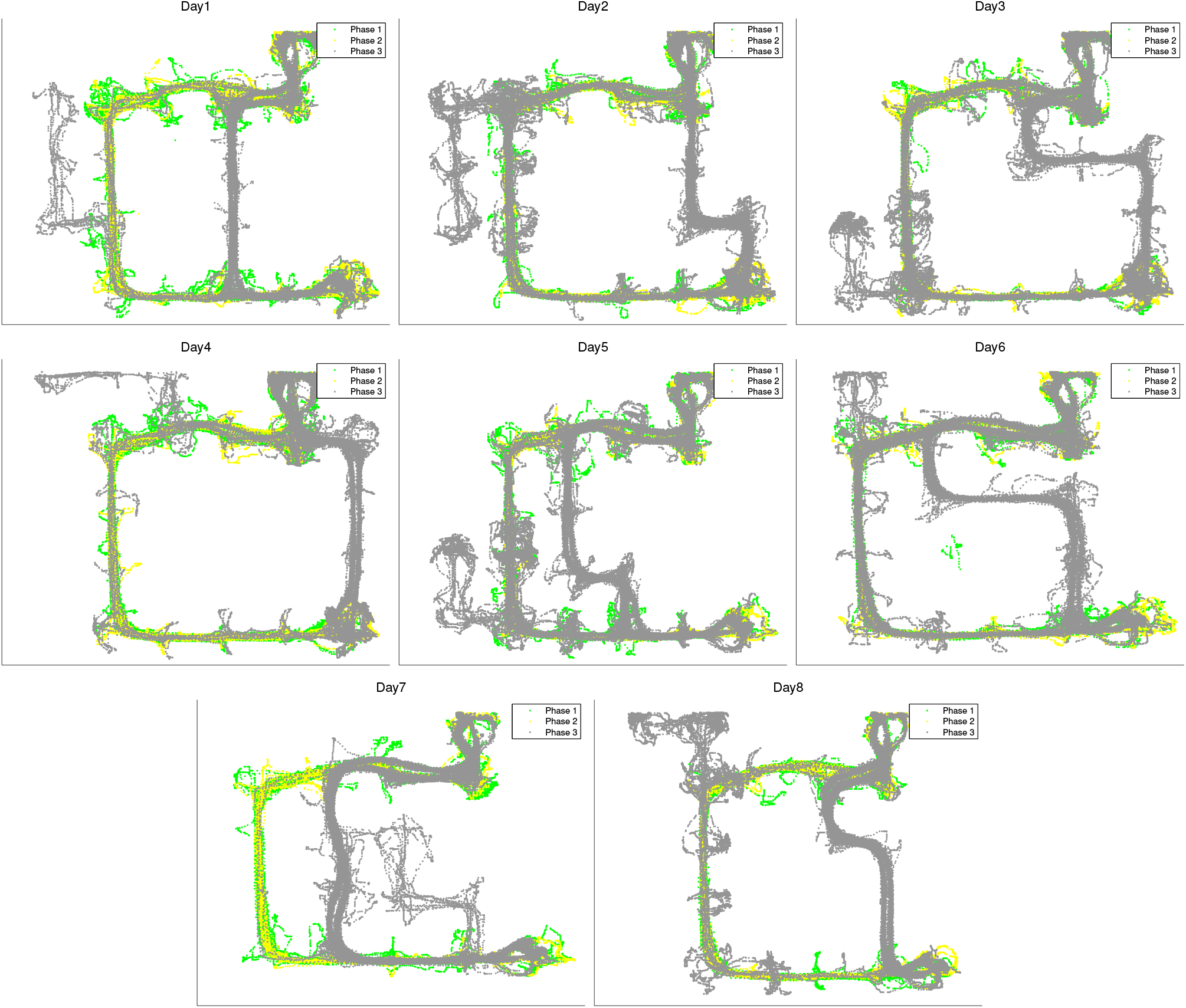
Dataset: trajectories of animal R01 across 8 days, 3 phases every day. See the main text for explanation.

### Experimental results, animal R01

In all the analyses below, we focus on trajectory data of animal R01 for illustrative purposes; see the Supplementary Material for data on animals R02 and R03, which show the same pattern of results as animal R01.

#### Building the dictionary of Primitives using sparse dictionary learning methods

The methodology to build and select the dictionary of primitives that we used in our subsequent analyses includes three main steps.

The first (preparation) step of our novel approach consists in dividing the database of rat trajectory data into three sets: a *training set*, which includes data from phases 1, 2 and 3 of day 1; a *validation set* and a *test set*, both including (non-overlapping) data from phase 3 of days 1 to 8. The actual data that we consider in the training, validation and test sets consist in “patches” of rat trajectories (i.e., portions of trajectories in x,y), lasting a maximum of 2.5 seconds each. Each of the three sets consists in 10000, non-overlapping patches, which are randomly selected. Patches with missing data (e.g., cases in which the animal goes out of the video camera frame) were excluded. Thus, the training data consist of an input matrix of dimension 10000×100, corresponding to 10000 patches of size 100 each composed of juxtaposed x,y position variables - (*x*(1), *y*(1),…, *x*(50), *y*(50)). Note that the time bin between two samples *x*(*k* + 1) and *x*(*k*) is fixed and is 0.05 *s*.

The complexity of these trajectory data can be appreciated by considering that a principal component analysis (PCA) shows that 73 principal components are necessary to explain about 95% of their variance, see Figure 5.

**Figure 5.**
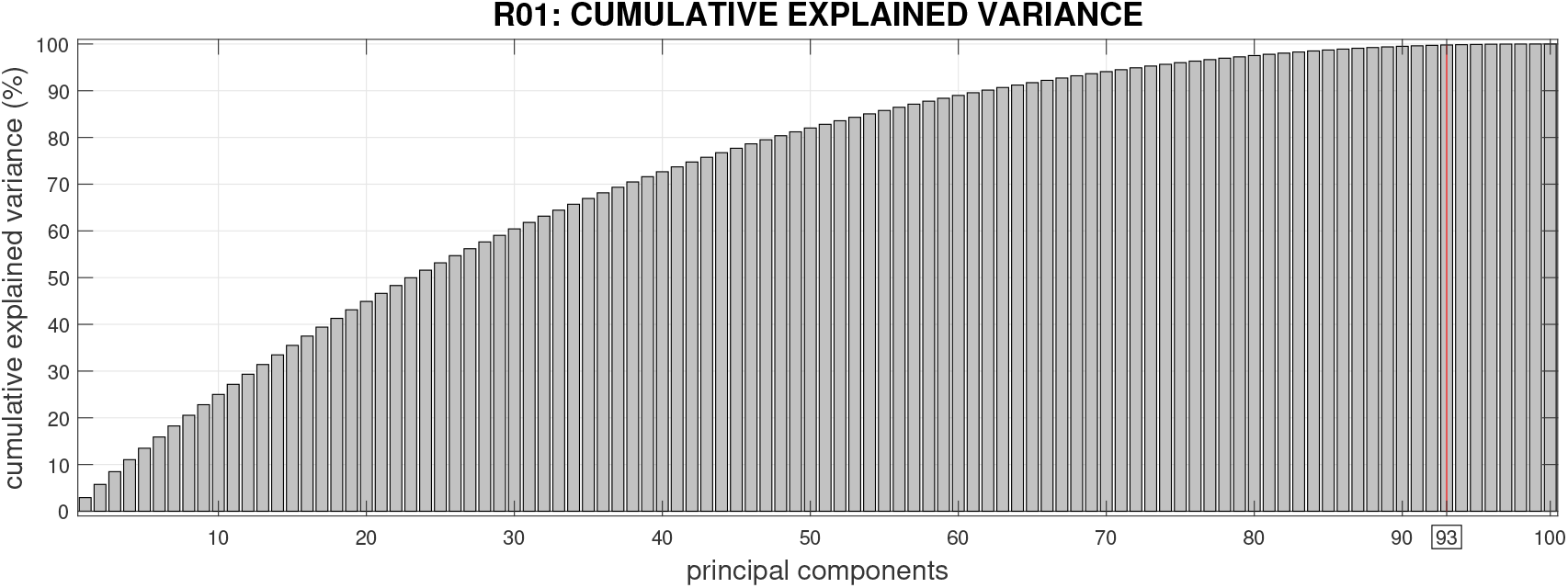
Cumulative explained variance in function of principal components. 93 principal components were selected in the tests, which explain about the 99% of the variance of the training set.

The second (learning) step of our novel approach consists in deriving candidate dictionaries of motor primitives which reconstruct accurately the *training set* data. For this, we use the Sparse Representation of Structured Sparse Dictionaries (SRSSD) approach, which directly implement our double sparsity hypothesis. SRSSD has two parameters of sparsity: coefficient and atom structured sparsity, which correspond to the two hypotheses of sparse superposition and sparse activity, respectively. The method therefore outputs not just dictionaries of motor primitives (*V*) but also their coefficients *U* (see Algorithm 1 and Methods Section for more details).

In this learning step, we select the best 600 (6×10×10) candidate dictionaries, obtained by varying 6 different dictionary sizes (with 25, 50, 75, 100, 125, and 150 motor primitives, respectively), 10 levels of atom sparsity and 10 levels of coefficient sparsity.

To measure how well the 600 candidate SRSSD dictionaries reconstructed the *training set* data, we adopted a methodology that is analogous to the *missing pixel method*: we canceled out some parts (from 10% to 90%) of the trajectories that compose the training set and evaluated how well the learned dictionaries permit to reconstruct these missing parts (see the Methods section for an explanation of how the reconstruction error RMS is calculated). Figure 6(left) shows coefficient sparsity and atom sparsity of the solutions found by SRSSD for dictionaries of size 150, i.e., the size that gave the best results, during the learning phase. The results show that the solution space is extensively covered, both in coefficient sparsity and in atom sparsity.

**Figure 6.**
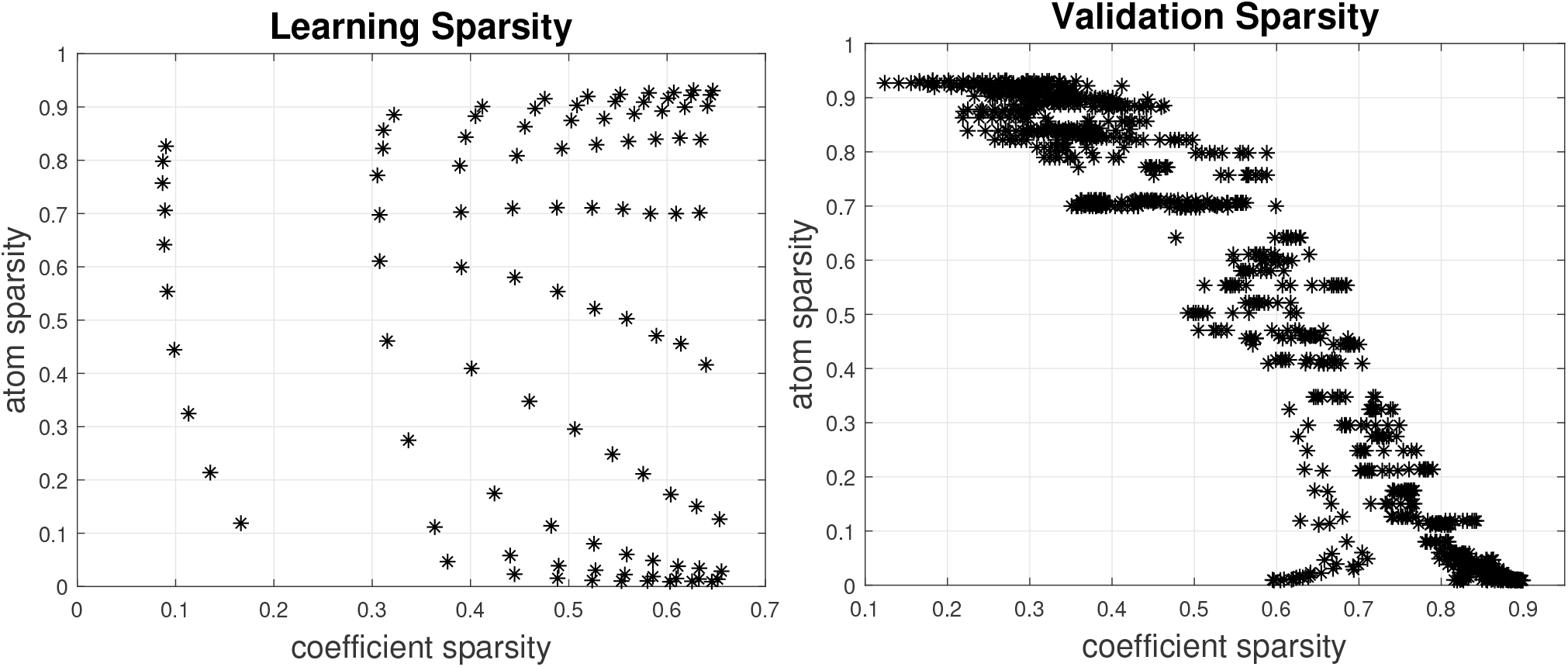
SRSSD method applied to navigation data from animal R01. Left: learning phase. Right: validation phase. Results are shown for dictionaries of size 150 (corresponding to lowest RMS in the test set / validation phase), with 10 values of coefficient sparsity and 5 levels of noise (0.1, 0.3, 0.5, 0.7, 0.9). Notice that the solution space is extensively covered, across both (coefficient and atom sparsity) dimensions.

In the third (validation) step of the approach we select the best 30 SRSSD dictionaries *V* amongst the aforementioned 600 candidates. For this, we force a reconstruction with sparse coefficients (see Algortihm 2): we fix *V* and obtain the couple *V* and *U* that better reconstructs the *validation set* data, by varying 5 levels of “noise” and 10 levels of coefficient sparsity.

While we conducted the validation on all the 600 dictionaries considered above, we only show the results of the 100 dictionaries of size 150 (i.e., the size that gave the best results during the learning step). Figure 6(right) shows coefficient and atom sparsity of the solutions in the validation phase of these 100 dictionaries, with 10 levels of coefficient sparsity and 5 levels of “noise” (0.1, 0.3, 0.5, 0.7 and 0.9) – corresponding to the percentage of “missing” trajectory data (10%, 30%, 50%, 70% and 90%) to be reconstructed. The results show the existence of solutions with both high coefficient sparsity and high atom sparsity, hence satisfying simultaneously the sparse superposition and the sparse activity hypotheses – and confirming that the range of the parameter space is well chosen. In the subsequent tests, we verify that successful solutions (those with better reconstruction) do satisfy a reasonably high double sparsity, hence lending support for both the sparse superposition and the sparse activity hypotheses.

The results of the validation step are shown in Figure 7 (black asterisks correspond to different dictionaries) and compared with PCA (horizontal line). The PCA results were computed by comparing the reconstruction error with 93 principal components explaining ≈ 99% of the variance of training data. The number of components (from 1 to 100) was selected by taking the best performing set that reconstructs the training set. The comparison illustrates that for each level of noise, various dictionaries have lower reconstruction error (RMS) than the best PCA subset. We also conducted a more systematic evaluation, in which we compared our approach based on double sparsity (SRSSD) with a method that entails no sparsity (PCA) and two other dictionary learning methods, each incorporating only one kind of sparsity: namely, coefficient sparsity *ℓ*_1_ *regularization*^34^ and atom structured sparsity SSPCA^35^. We compared the best instances of SRSSD, SSPCA, and *ℓ*_1_, which resulted from considering 10 different values of sparsity parameters in the same range (coefficient sparsity and atom sparsity parameters for SRSSD, coefficients sparsity parameter for *ℓ*_1_ and atom sparsity for SSPCA) and best PCA set of 93 components, as described above. Each method can be seen as the computational counterpart of a different biological hypotheses: no sparsity (PCA), sparse activity (SSPCA), sparse combination (*ℓ*_1_) and double sparsity (SRSSD).

**Figure 7.**
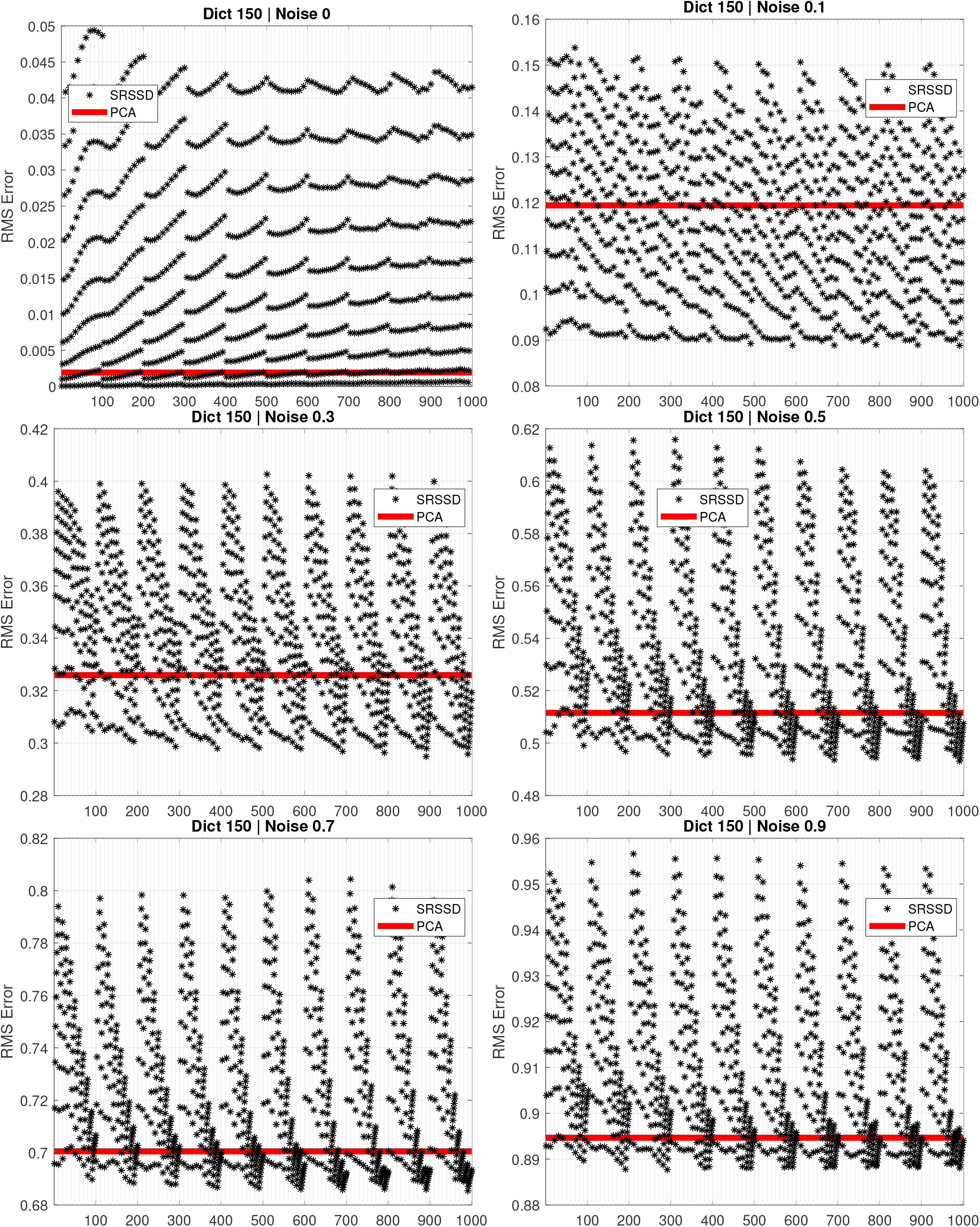
Comparison between the reconstruction (RMS) error of SRSSD (blue dots) and PCA (red line) across various levels of noise. Each figure shows 100 dictionaries found during the learning step. We remind that each reconstruction relies on 2 kinds of sparsity: atom structured sparsity (varied during the learning phase) and coefficient sparsity (which is varied here); see the Methods section. Here, each of the 100 dictionaries (with its fixed atom sparsity) is tested with 10 different values of the coefficient sparsity parameter. Hence, for each figure, each of the 1000 asterisks corresponds to a unique combination of dictionary and coefficient sparsity. The reconstruction error for all the asterisks that appear under the red line is better than PCA.

In this comparison, we also considered another method, *ℓ*_1_-RD, which starts from a random dictionary with values in the interval (1, 1) matching the dimensionality of the other sparse methods (SRSSD, SSPCA, *ℓ*_1_). We generated 1000 random dictionaries and selected the best one, which was capable of reconstructing the training set with a sparse reconstruction, varying the sparse coefficient parameter in a similar way as *ℓ*_1_ method, but without optimising the dictionary values. This method allows us to experimentally evaluate the possibility of obtaining a useful dictionary representation by chance.

We compared the five methods with 6 levels of noise (0%, 10%. 30%, 50%, 70%, 90%). For each level of noise, we computed the percent error of the methods in relation to SRSSD, as follows:

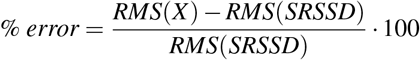

where *RMS*(*X*) is the best reconstruction error of a selected method *X* and *RMS*(SRSSD) is the best reconstruction error of SRSSD.

The results of this comparison are shown in Figure 8 (note that positive values indicate smaller reconstruction error). Our results show that, as expected, higher noise levels decrease reconstruction accuracy across all methods. Moreover, and most importantly, SRSSD outperforms all the other methods across all levels of noise and is more noise-tolerant.

**Figure 8.**
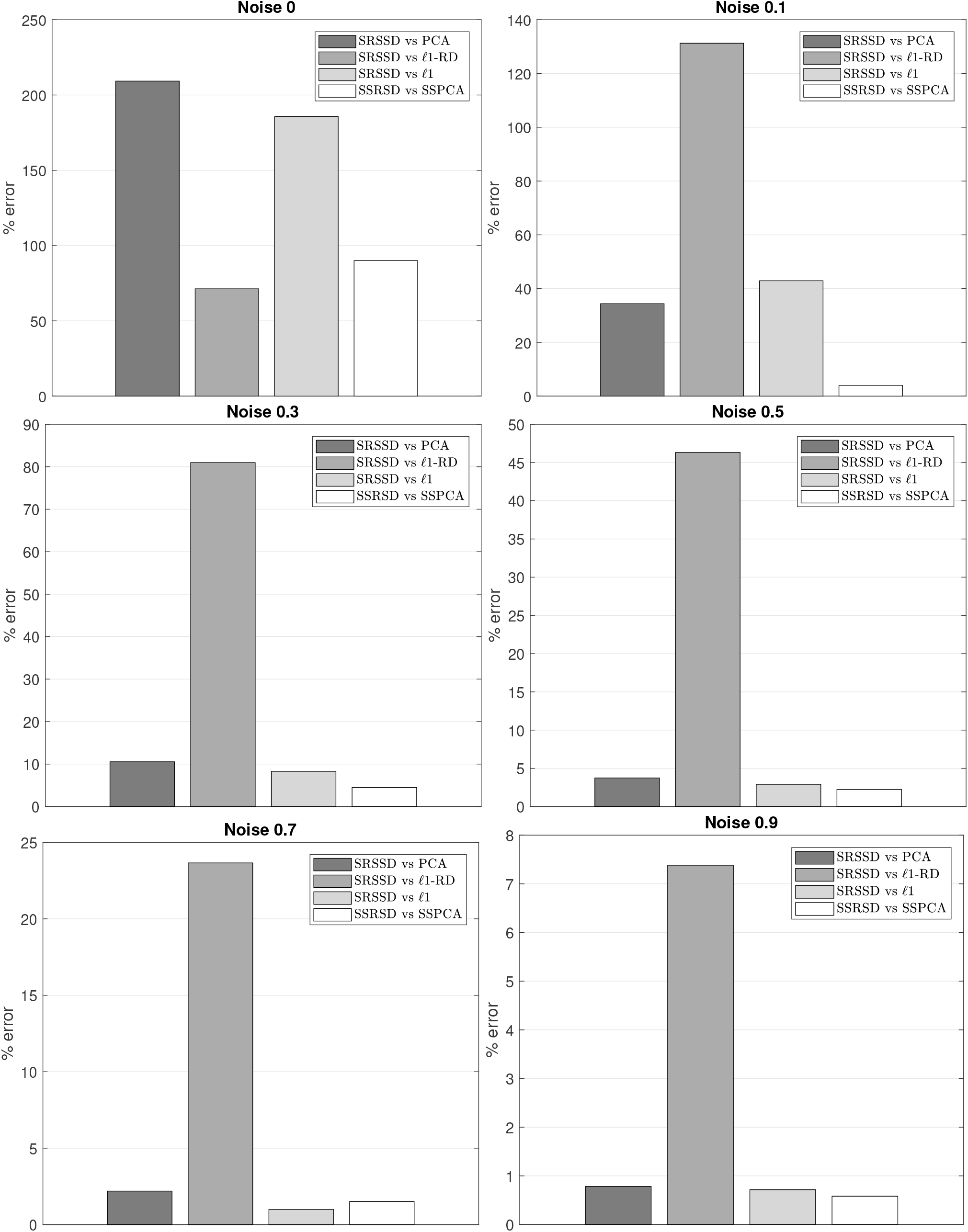
The percent error for PCA, *ℓ*1-RD, *ℓ*1-regularization, and SSPCA compared to SRSSD, across six levels of noise (0%, 10%. 30%, 50%, 70%, 90%). It is possible to appreciate that SRSSD outperforms the other methods, across all noise levels.

In sum, this first series of tests and comparisons indicates that the choice of a double-sparse approach like SRSSD nicely incorporates the double sparsity hypothesis, which is appealing from a *biological* perspective; and furthermore, it affords a more efficient reconstruction and prediction of motor data of the rat, which is appealing from a *computational* perspective.

Consequently, in the subsequent sections, we shift our focus on a deeper analysis of the double sparsity method by means of the best 30 SRSSD dictionaries learned in these tests and illustrate the potentialities of the proposed approach.

#### Reconstruction across consecutive days using the best 30 dictionaries

We tested whether the best 30 dictionaries *V* identified using the above procedure afford an accurate reconstruction of novel data, from the *test set*. More specifically, we were interested in assessing whether a learning procedure that only considered data from day 1 (i.e., *training set*) permitted reconstructing non-overlapping data from the third phases of days 1-8 in our dataset (i.e., *test set*). Note that the reason why the *test set* only includes the third phases of each day is that this is when, in the rodent experiment, novel portions of the maze appear. Data on the third phases of each day are this ideal to test whether our approach generalises to novel and unforeseen situations. Furthermore, we asked whether the motor primitives required to reconstruct animal trajectories in a given day capture essential characteristics of the day’s task, such as maze complexity or stereotypy of behavior.

Figure 9 shows the reconstruction results for the best 30 SRSSD dictionaries. There are at least two elements to be appreciated. First, reconstruction error is lower in days 1, 2 and 8. The low reconstruction error in day 8 indicates that the method generalises well across distal days. The fact that reconstruction is better in day 8 than the preceding days could be explained by considering the close spatial similarities between the mazes of days 1 and 8 (see see Figure 4). Second, as shown in Figure 9(b), days 4 and 8 have the highest coefficient sparsity, implying that fewer motor primitives are required for their correct reconstruction. The fact that behavior in different mazes and days can be reconstructed using small subsets of (possibly overlapping) primitives lends support for the biological sparse superposition hypothesis.

**Figure 9.**
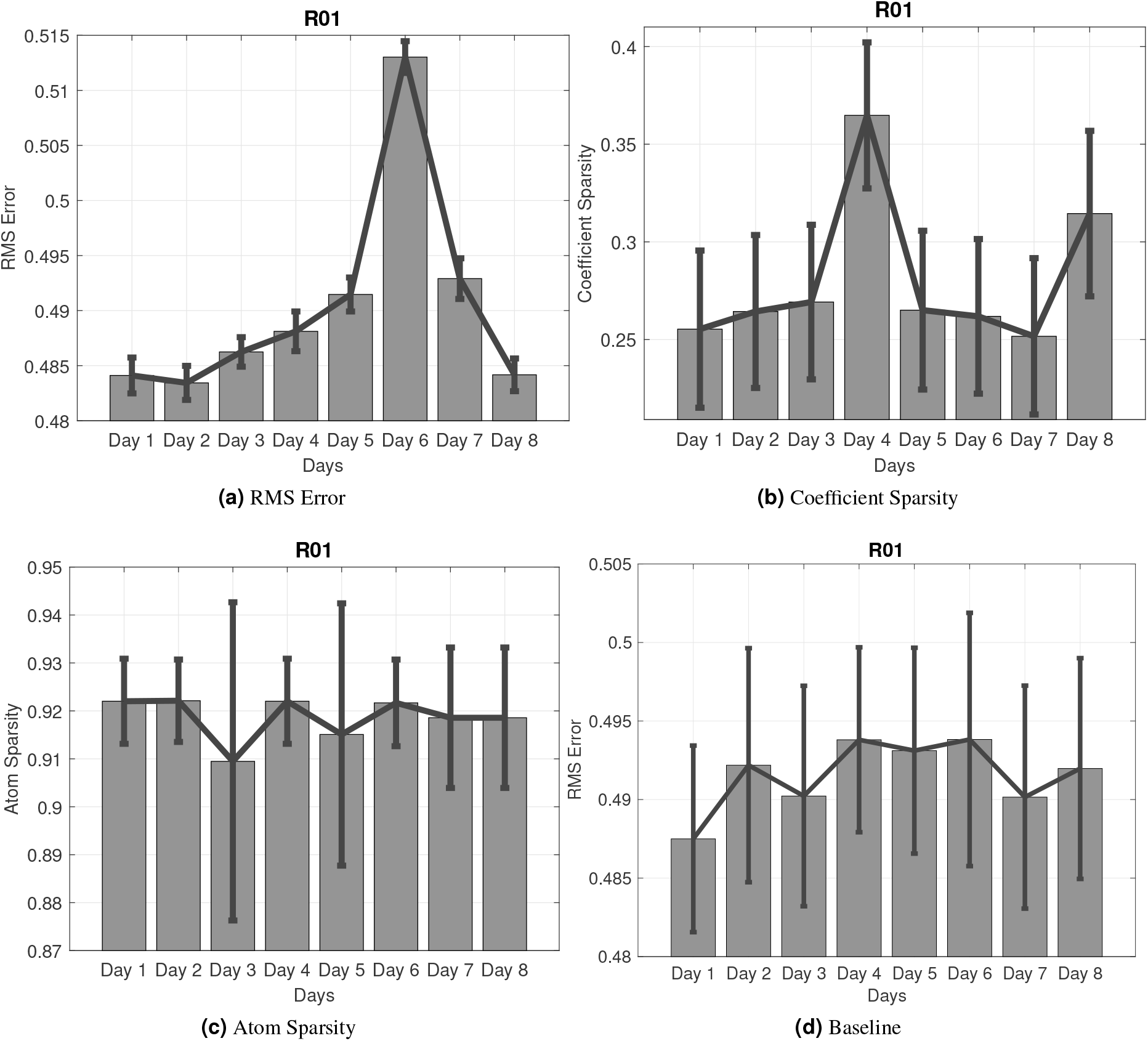
Reconstruction across 8 days (after learning only from data of day 1. Noise is 0.5 and the best 30 SRSSD dictionaries found on day 1 are used. Panel (a): RMS Error. Panel (b): coefficient sparsity. Panel (c) atom sparsity. Panel (d) Baseline performance: reconstruction of Phase 1. Note that Panel (a) shows an increase in the reconstruction error, until day 6 and then lower levels of the error in the next two days. The increase of reconstruction error across successive days is to be expected, given that the dictionary is acquired on day 1 (the high error value of day 6 may be due to specific characteristics of the corresponding maze). The decrease of error in the last two days may be due to stereotypy of behavior, specific characteristics of the corresponding mazes, or a combination of both factors.

Most importantly, we found the lower coefficient sparsity of days 4 and 8 to reflect well maze characteristics and animal behaviour. This can be appreciated if one consider our next analysis of task demands and complexity, see Figure 10. The first measure of task demands we considered is the mean time the animal needed to complete a trial, see Figure 10a. This analysis shows that time is shorter in days 4 and 8, which are the two days in which our previous analysis revealed higher coefficient sparsity, see Figure 9(b).

**Figure 10.**
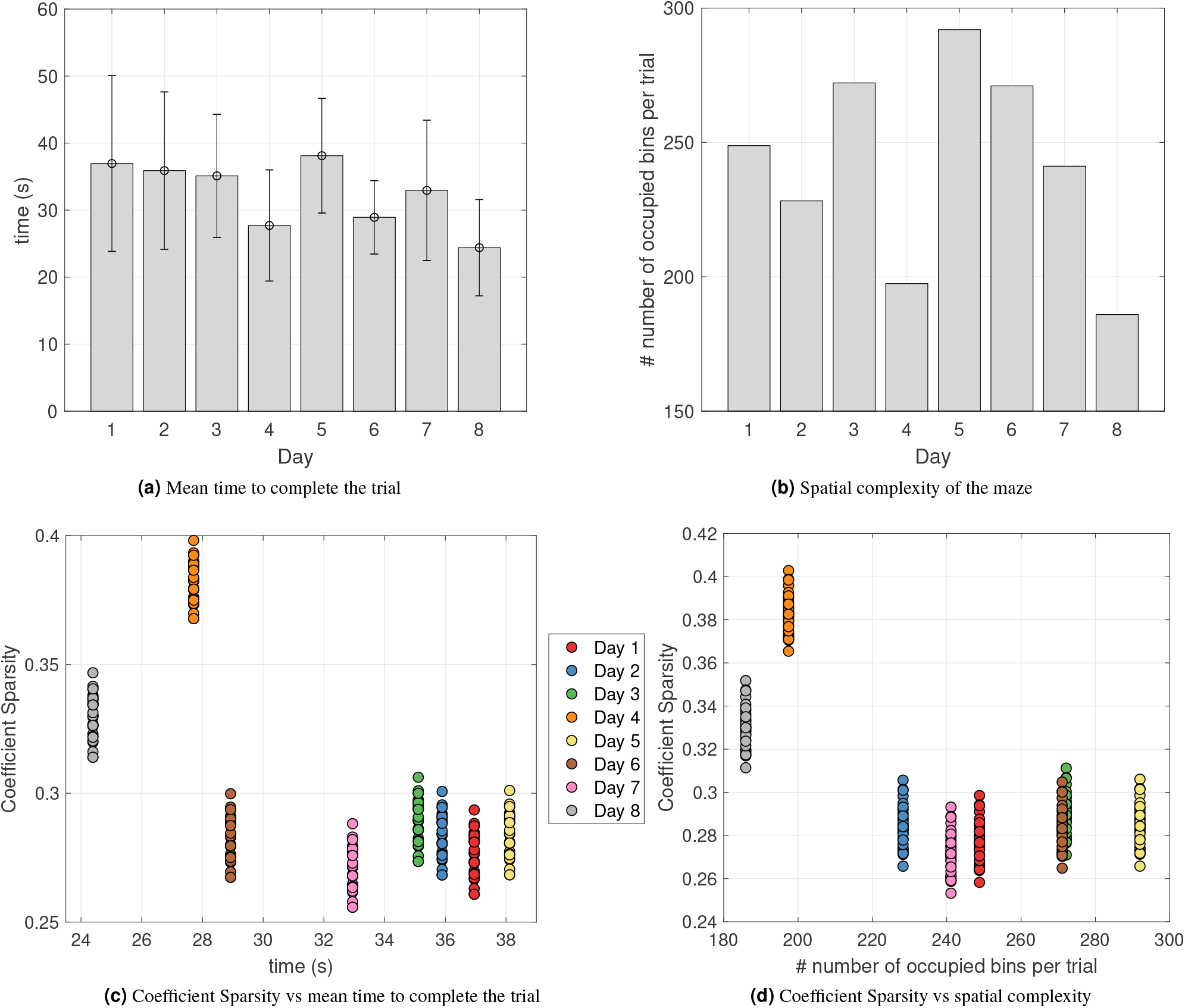
Measures of task demands and complexity of the rodent task. (a) Average time in seconds and standard deviation required to complete a trial and secure a reward. (b) Spatial complexity of the maze: number of bins per trial occupied during the task. (c) Scatter Plot of coefficient sparsity versus time to complete the trial. (d) Scatter Plot of coefficient sparsity versus spatial complexity.

The second measure we considered is the spatial complexity of the maze, see Figure 10b. We assessed spatial complexity by dividing the maze into bins, counting how many bins the animal occupies for each trial of phase 3, and then averaging (i.e., dividing this number by the number of trials); see the Supplementary Material for Figures illustrating the results of this process. High spatial complexity (or entropy) implies that the animal has explored and occupied all the available positions, while low spatial complexity means that the animal followed stereotyped trajectories. Note that there are multiple ways to construct spatial complexity^36,37^. In our approach, complexity depends on behavior (similar to^38^) – or better, on a graph constructed on the basis of the actual trajectories of the animal, rather than a fixed graph that only includes spatial dependencies. Our analyses reveals that spatial complexity of days 4 and 8 is lower compared to the other days, revealing stereotyped behavior in these days (see also below on a convergent measure of stereotypy). The same results emerge if we plot coefficient sparsity versus time to complete the trial (Figure 10c) and coefficient sparsity versus complexity (Figure 10d). Note that days are color-coded and days 4 and 8 (orange and grey, respectively) have the highest values in both measures.

Overall, this analysis reveals that in days 4 and 8, coefficient sparsity is higher, the animal is faster and maze complexity is lower. The coherence across these measures supports the idea that the motor primitive formalism we adopted captures essential task components such as its complexity.

### Example applications of the method in the generation, description and classification of behavior

Animal behavior is highly structured. Behavioral scientists and neuroscientists often deal with complex datasets of behavioral data – e.g., the full spatiotemporal x(t), y(t) data – that are not convenient for asking questions like: how stereotyped is the animal’s behavior? How well can future behavior be predicted by past behavior? How does behavior evolve with experience on the task, within and across sessions?

One appeal of the motor primitive code is that it provides a description of rat behavior that is more compact than the full spatiotemporal [x(t),y(t)] data yet richer than a simple mono-dimensional descriptor [e.g., speed(t)] – and lends itself to a number of applications. Our method based on motor primitives permits estimating the building blocks of a “computational ethogram” that captures the main behavioral regularities (i.e., repeated spatiotemporal sequences that are highly predictable within-sequence) and the temporal dependencies between them (e.g., transitions or contingencies between sequences). This method can be used in a number of ways.

#### Generating simulated behavior on novel mazes and associated neuronal responses

One possible application would be generating simulated behavior on various novel mazes. Simulated trajectories can be helpful in estimating the to-be-expected complexity of behavioral data. Furthermore, they can be fed as input to other components to predict hippocampal place cell activity, thus affording power analyses. One example is shown in Figure 11. We used the computational models of^39,40^ to simulate the firing of example grid and place cells during uniform exploration (Panels (e) and (f), respectively) versus partial exploration of the environment following a simulated trajectory (Panels (c) and (d), respectively), also inferring the connectivity structure that link the simulated place cells (S1 to S20 in Panel (g)) This example illustrates the possibility to use simulated trajectories to predict neuronal firings in novel environments. Note that in the above examples, the trajectories are generated by chaining motor primitives randomly, mimicking uniform exploration. Future studies may consider extensions of this method that learn (flat, hierarchical or context-dependent) transition probabilities between primitives and can generate more contextually-adequate trajectories.

**Figure 11.**
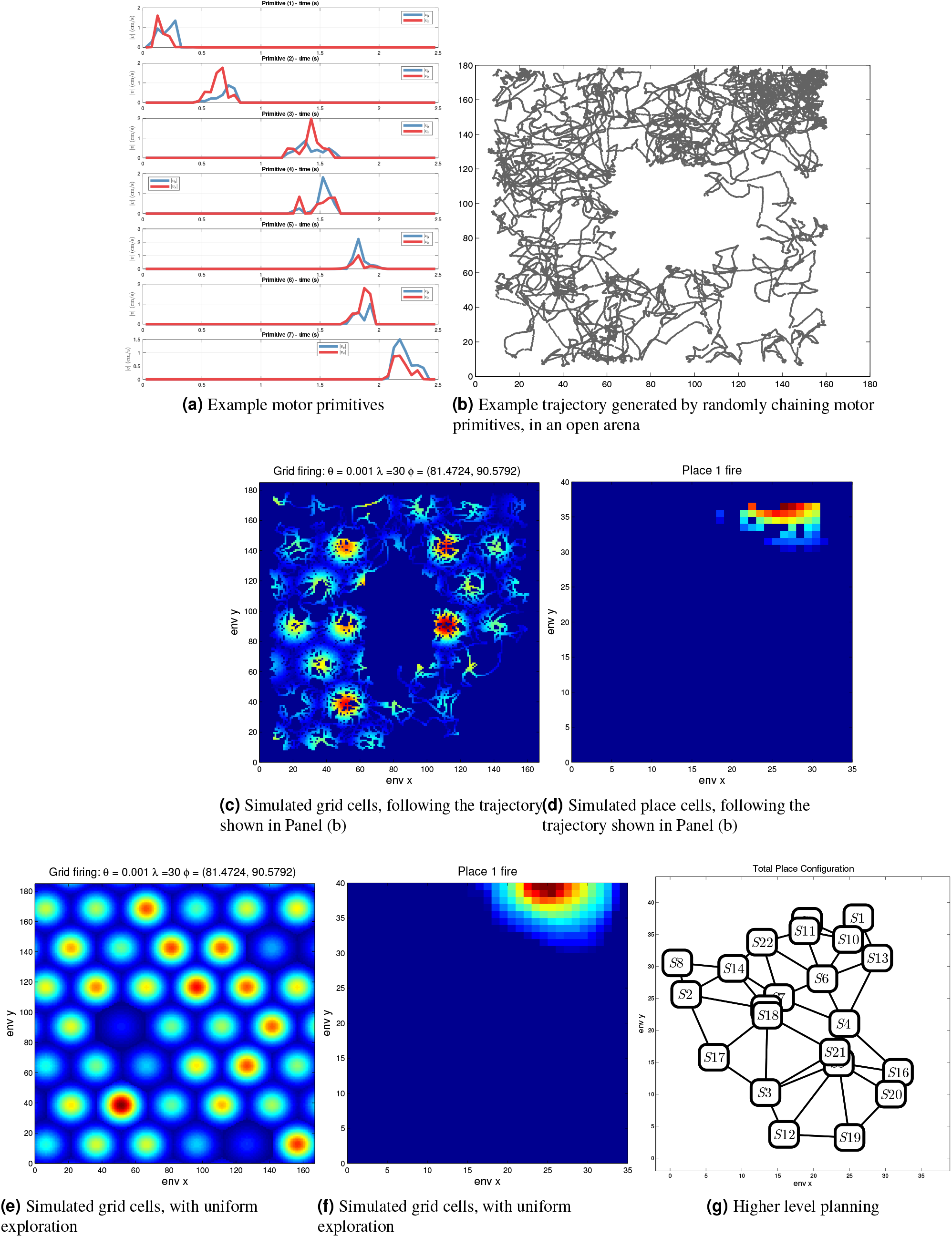
Graphical illustration of potential applications of the analysis. Panel (a) provides examples of motor primitives extracted from a maze, which could be used to simulate animal behavior in novel mazes, such as the open arena shown in Panel (b). In turn, the simulated trajectories shown in Panel (b) can be used as inputs to computational models that permit simulating grid cells (Panel (c)) and place cells (Panel (d)); which are different from grid cells (Panel (e)) and place cells (Panel (f)) generated under the hypothesis of uniform exploration. It is also possible to infer the connectivity structure that link the simulated place cells (S1 to S20 in Panel (g)), by considering which ones are linked via motor primitives. In principle, this connectivity structure can be used to design planning algorithms.

#### Assessing behavioral stereotypy after learning

Another application is testing stereotypy of behavior (as a possible index of habitization), by looking at coefficient sparsity during learning. Stereotypy of behavior can be tested by considering whether coefficient sparsity increases during learning, indexing the fact that the animal is using fewer primitives (and hence a more restricted and stereotyped behavioral repertoire). As an example of this analysis, we considered how coefficient sparsity changes while animal R01 learns to navigate the basic U-shaped maze (i.e., during phase 1 of day 1). For this, we have split data of phase 1 of day 1 into 5 intervals of the same length, removed the two extremes (1 and 5) and the middle interval (3), and compared intervals 2 vs. 4, corresponding to initial and final phases of learning, see Figure 12. One-way analysis of variance (ANOVA) indicates that the difference between coefficient sparsity values of intervals 2 and 4 is statistically significant (*F*[1,78] = 4.197, *p* < 0.05), with the latter interval showing greater coefficient sparsity. These results indicate that as the animal R01 learns the task, it uses less primitives and a more stereotyped behavior. Assessing whether this method is potentially more robust than measures of path stereotypy^20,41^ or simple descriptions of x(t), y(t) data like Fourier decomposition^42^ is beyond the scope of this work. Yet, in general, describing spatiotemporal data in terms of (movement) primitives is considered to be more noise-tolerant in motor control^1–4^.

**Figure 12.**
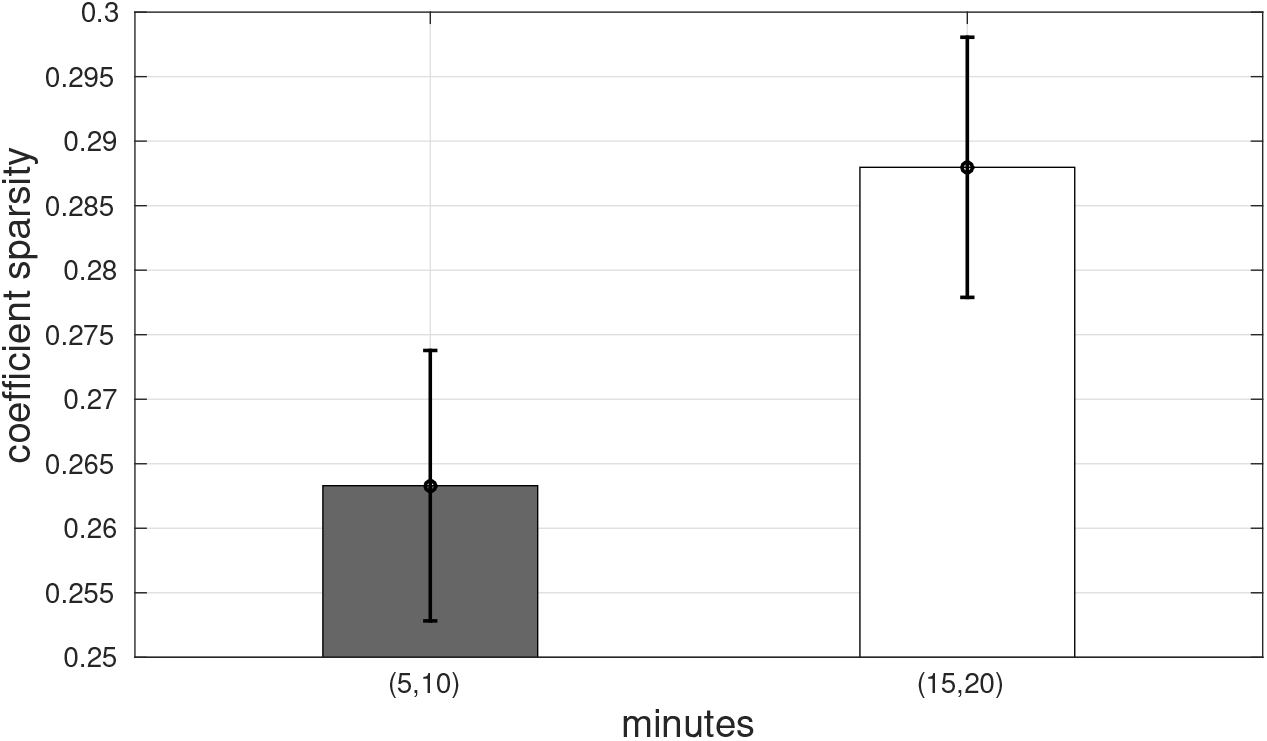
Mean and standard error of coefficient sparsity values for animal R01 behavior in the 2nd and 4th intervals of phase 1 of day 1. The coefficient sparsity values relative to the animal behavior range in [0.25, 0.30]. While the animal learns to navigate in the U maze, coefficient sparsity increases significantly (*ℓ*[1,78] = 4.197, *p* < 0.05), revealing an increased stereotypy of behavior.

#### Classifying and predicting behavior

Our approach based on the identification of motor primitives can also be used to classify and predict animal behavior and choices. To exemplify this, we performed an experiment aimed at studying whether motor primitives extracted during different trials of phases 3 of each day permit to classify whether in these trials animal R01 follows the U-path or the shortcut. For each day, we randomly select 1000 trajectories (500 Shortcut and 500 U-path trajectories) and apply the reconstruction method, for each of the 30 best dictionaries, in order to find sparse coefficients. We then use the coefficients to classify U-path versus shortcut trajectories (which we had previously labelled, see examples in the Supplementary Materials). For this, we perform a classification test on the set of 1000 trajectories, using a linear support vector machine (SVM) classifier in a k-fold cross validation (with k=5). We compute mean and standard deviation over the dictionaries of the accuracies. Figure 13 shows that this method affords very accurate classification across all the 8 days. Note that this result does not trivially depend on the fact that the animal occupies different x-y coordinates during U-path versus shortcut trajectories. The motor primitives are agnostic about maze-centered x-y coordinates, as their reference system is centered on the animal, see Figure 1a.

**Figure 13.**
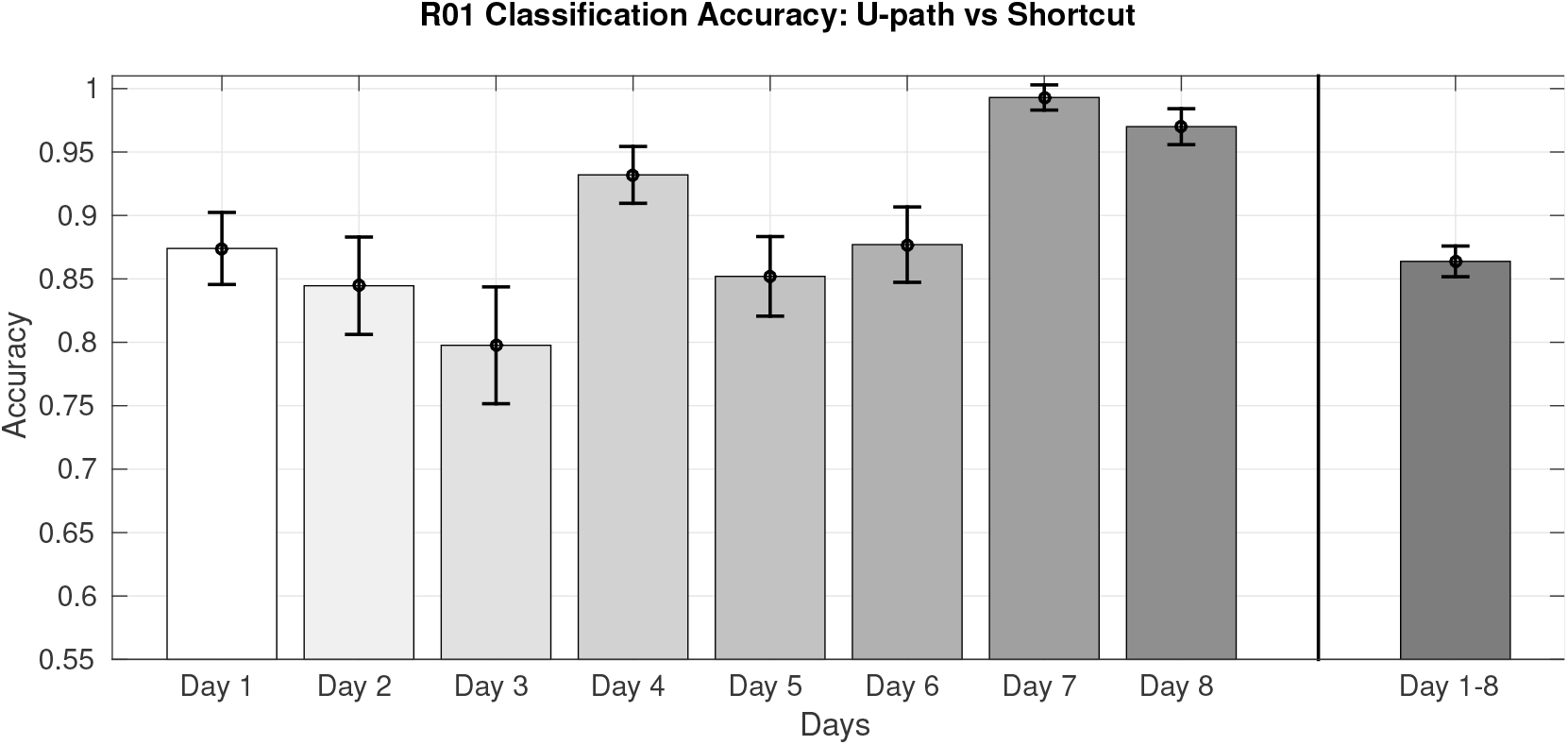
Classification of trajectories (U-path or shortcut) of phase 3, animal R01s

The last bar of Figure 13 (labelled Day1-8) shows the results of a second experiment, in which we used a single dataset of 8000 samples of trajectories (1000 for each day, resulting in 4000 U-path and 4000 shortcut trajectories, combined together). Note that while the U-path remains constant across days, the shortcuts change every day, making this experiment potentially more challenging than the former. Again, we performed a k-fold cross validation test using a linear SVM classifier and we computed accuracies over the 30 dictionaries coefficients. Even in this second experiment, our method affords very accurate classification of U-path versus shortcut trajectories. These results illustrate that motor primitives extracted through our method capture behaviorally relevant regularities that are informative (for example) about animal trajectories and choices.

## Discussion

We introduced a novel framework based on sparse dictionary learning to quantitatively characterize rodent behavior during spatial navigation. This approach is capable of extracting latent structure – a “statistical ethogram” – from rodent motor data (i.e., position and velocity), in terms of movement building blocks of motor primitives. The approach adopted here (SRSSD) specifically incorporates the hypothesis that motor primitives are organized in terms of two sparsity principles: each movement is controlled using a limited subset of motor primitives (sparse superposition) and each primitive is active only for time-limited, time-contiguous portions of movements (sparse activity).

Our experiments on rodent trajectories in eight mazes show that the double sparse dictionary learning (SRSSD) method outperforms principal component analysis (PCA) and two other sparse dictionary learning methods, which incorporate only sparse coefficients (*ℓ*_1_ -*regularization*) or atom structured sparsity (SSPCA), across all the noise levels we tested, with appreciable results over 0.5 noise – thus speaking in favour of the robustness of the methods. Furthermore, our results show good generalization across eight consecutive days. Despite the fact that we only used data from day 1 in the training set, the reconstruction of trajectories was not degraded in the successive seven days, suggesting that the motor primitives approach extracts regularities that are persistent over time. These regularities correspond to spatiotemporal patterns of movement, which can include, for example, rapid forward movements or rotations, which are repeated multiple times during the maze navigation and may be thus revelatory of an underlying modular organisation of behavior.

Our further analysis indicates that our approach based on motor primitives characterizes well task regularities such as spatial complexity and movement variance in terms of (coefficient) sparsity constraints. This makes intuitive sense, as navigation scenarios in which the animals’ movements had lesser variance (e.g., for days 4 and 8 from animal R01) require less motor primitives (i.e., allow for more sparsity). It is also reassuring to notice that, although our method allowed the construction of motor primitives of up to 10 second, the primitives that actually populate the best dictionaries are of about 1 second, which is a more realistic time constraint for biological movement.

This is, to the best of our knowledge, the first framework that applies the concept of motor primitives – which is popular in motor control and computational neuroscience – to a data-driven analysis of rodent spatial navigation data. The method introduced here is based on 2D position data, which could be seen as limited compared to more sophisticated (3D camera, accelerometer) approaches^21^. However, functionality with 2D data is more applicable to the vast majority of behavioral scientists and neuroscientists.

The approach we have devised can be used by behavioral scientists and neuroscientists in behavioral and neural analyses. Some example uses that we have shortly illustrated in this paper are: controlling for confounding effects (e.g., of maze complexity on behavior and reward collection), analyzing habitual or stereotyped behavior, classifying and predicting animal choices, and predicting place and grid cell displacement in novel mazes. The results we reported illustrate the feasibility of the method, which can be successfully added to the toolbox of behavioral scientists and neuroscientists to improve their ability to quantitatively characterize and understand animal behavior.

Besides, our approach can be used for other purposes, such as, for example, to support the analysis of neuronal data or to design experiments that test the neuronal underpinning of motor primitives for spatial navigation. In decoding analyses, tuning curves (aka, the encoding model^43^) are typically estimated from those epochs during which the animal is running, which in turn is estimated using a running speed cutoff. However, the cutoff cannot clearly distinguish between different situations, e.g., grooming, directed running towards a goal or performing exploratory moves. Distinguishing these different behaviors is important as they modulate neural firings, above and beyond spatial position. The approach presented here can distinguish different movement primitives (or even behavioral patterns or modes) and has thus the potential to improve our ability to relate neural activity to behavior, as done e.g. in decoding analyses. Furthermore, the motor primitives extracted from behavior may be characterized neurally. Studying the potential neuronal correlates of (spatial) motor primitives remains an important objective for future research.

Finally, our approach can be extended to reveal and study more complex patterns of behavior than those considered here. Note that the method we have described does not try to estimate sequential dependencies between primitives. However, it would be trivial to compute those dependencies (e.g., a probability distribution *P*(*pi|p j*) that a primitive pi follows a primitive pj) from training data, in order to estimate the most likely (or most surprising) transitions. It would be then possible to derive measures of path novelty, which would be related to (relatively) unpredictable transitions; or to study sequential (possibly, planning-related) patterns of behavior. These and other potential applications of the method remain to be tested in future studies.

## Materials and methods

To characterize and extract motor primitives from trajectory data (i.e., position and velocity) during spatial navigation we adopt *sparse dictionary learning* techniques ^31–33,44–46^. Sparse dictionary learning techniques consist of unsupervised learning of a set of atoms that can be linearly superimposed to represent the elements of the dataset (in our case rodent spatial trajectories). In these approaches, either the atoms or the coefficients of the linear combinations (or both) are imposed to have a large number of components equal to zero. These two cases correspond to *atom structured sparsity* and *coefficient sparsity*, respectively. In dictionary learning literature a reconstruction that forces sparse coefficients is also referred to as *sparse coding*.

These two kinds of sparsity distinguish dictionary learning methods from more traditional approaches like PCA, which decompose data using orthogonal basis functions. Sparse representations are instead based on the construction of a set of nonorthogonal – *over-complete* – representations, usually called *dictionaries*; see^47^ for a thorough review.

Under an over-complete basis, the decomposition of a signal is not unique, but this can offer clear advantages. Firstly, it results in a greater flexibility in capturing structure in the data: instead of a small set of general basis functions, there is a larger set of more specialized basis functions. In this way, relatively few elements are required to represent parts of the learning signals. This eventually results in more compact representations, because each basis function can describe a significant amount of structure in the data. In pattern recognition literature, these over-complete dictionaries are able to achieve more meaningful and stable representations of the source data^48^. For this reason, sparse methods generally outperform the traditional orthogonal basis functions^49^. Another advantage of dictionary learning methods over PCA is that the length of the primitives does not need to be predefined or fixed, but it is automatically determined by the algorithm, allowing for more flexibility depending on the particular application case.

### Dictionary learning and the SRSSD approach

Recent machine learning approaches^44–46,50–53^ represent signals as linear combinations of a large number of *atoms*, collected in sets called *dictionaries*. The dictionaries are computed using prior information encoded in penalization terms defining a particular kind of a minimization problem. These approaches are informatively grouped into two classes:

i. *Sparse atoms*. This is the case when each atom involves just a small number of the original variables (see, for example, Sparse-PCA^50^, sPCA-rSVD^54^ and Structured-Sparse-PCA^35^);
ii. *Sparse coding*. In this case, an *overcomplete* set of atoms is learned from the data, but the approximation of each signal involves only a restricted number of atoms. Hence, signals are represented by sparse linear combinations of the atoms (see, for example, MOD^55^, *K*-*SVD*^53^, and *ℓ*_1_-regularized^34^).

Recently, a new algorithm named Structured Sparse Dictionaries for Sparse Representation (*SRSSD*) was presented in^33^ and applied to a cognitive science study^32^; its fundamental characteristics is to search for sparse atoms and sparse coding simultaneously, reconciling, in this way, the two different dictionary-learning approaches.

Within the SRSSD method, we denote **X** ∈ ℝ^*n*×*p*^ as a matrix where rows correspond to experimental observations (e.g., sequences of x and y coordinates that compose a trajectory, e.g., x1, y1, x2, y2, etc.). Note that *n* is the number of patches and *p* their length. Furthermore, we denote **V** ∈ ℝ^*p*×*r*^ as a *dictionary*, whose *r* columns **V**^*k*^ represent the atoms learned by the dictionary. Thus, *r* is the number of atoms and *p* their maximum length. Note that given its sparsity, the SRSSD method can learn dictionary elements having different length, whose maximum we set to *p* (e.g., the number of nonzero consecutive elements can be < *p*). Finally, we define **U** ∈ ℝ^*n*×*r*^ as the coefficient matrix.

The aim of SRSSD is finding out the best approximation of **X** in terms of **V** and **U**(i.e., **X** ≈ **UV**^T^). This problem can be formulated in terms of a minimization problem as follows:

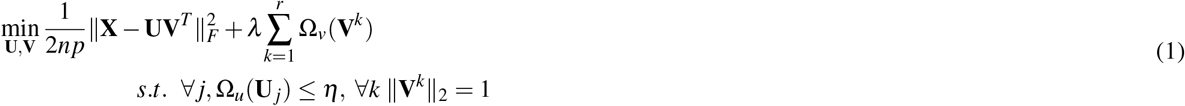

where _*F*_ is the Frobenius norm for matrices, **U**_*j*_ are the rows of **U**, Ω_*v*_(**V**^*k*^) and Ω_*u*_(**U**_*j*_) are norms or quasi-norms constraining (*regularizing*) the solutions of the minimization problem, with the parameters *λ* ≥ 0 and *η* ≥ 0 that control to which extension the dictionary and the coefficients are regularized, respectively. If one assumes that both Ω_*u*_ and Ω_*v*_ are convex, the problem (1) is convex w.r.t. **U** for **V** fixed, and vice versa.

Following^35^, the atom structured sparsity is imposed by setting

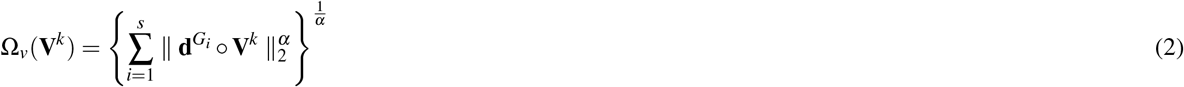

where *α* ∈ (0, 1), and each 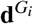 is a *p*-dimensional vector satisfying the condition 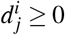, with 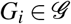 where 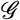 is a subset of the power set of {1,…, *p*}, such that 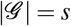 and 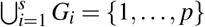. Thus, the vectors **d**^*i*^ define the structure of the atoms. More specifically, each 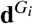 individuates a group of variables such that 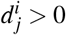 i f *j* ∈ *G_i_* and 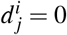 otherwise. The norm 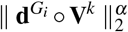 penalizes the variables selected by 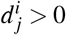, hence, by this norm, each vector 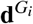 induces non zero values for the *j*-th elements of the atom **V**^*k*^, when *j* ∈ *G*.

The resulting set of selected variables depends on the contribution of each 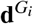 as described in^56^. For example, if the vectors 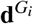 represent a partition on the set {1,…, *p*}, then the penalization term (2) favours atoms **V**^*k*^ composed of non-zero variables belonging to just one part of the partition and so on: for specific choices of 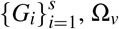 leads to standard sparsity-inducing norms.

Nevertheless, the norm Ω_*v*_ expressed in Equation (2) is not differentiable and, consequently Equation (1) is not convex with respect to **V** for **U** fixed — although the converse is still true. By using results presented in^35,57^, we can write down Ω_*v*_ as a quadratic expression and reformulate the Equation (1) as:

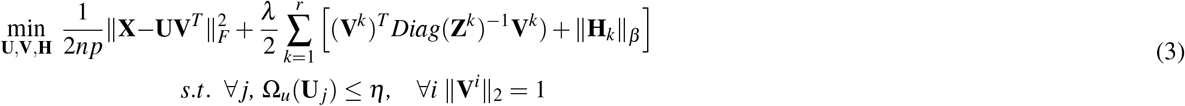

where 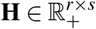 is a matrix satisfying the condition *h_ki_* ≥ 0 with 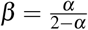 and that minimizes the second term of the previous expression in which **Z** ∈ ℝ^*p*×*r*^ has got its own elements defined as 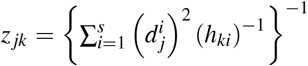. Notice that, as shown in^35^, for both **U** and **V** fixed, the minimizer of (3) can be given in the closed form 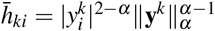, for *k* = 1, 2,…, *r* and *i* = 1, 2,…, *s*, where each **y***^k^* ∈ ℝ^1×*s*^ is the vector **y***^k^* = (‖**d**^1^ ○ **V**^*k*^‖_2_, ‖**d**^2^ ○ **V**^*k*^‖_2_,…,**d**^*s*^ ○ **V**^*k*^‖_2_). Ultimately, we impose the constraint ‖**V**^*i*^‖_2_ = 1 to avoid solutions where **V** goes towards **0**, or **U** becomes a sparse matrix, whose non-zero elements have excessively high values.

Since the functional in (3) is separately convex in each variable, we solve the minimization problem following the usual approach of alternating optimizations with respect to the values **H**, to the coefficients **U** and to the dictionary **V**. Most methods are based on this alternating scheme of optimization^34,58^ and its convergence towards a critical point of the functional is guaranteed by Corollary 2 of^59^.

Algorithm 1 summarises the procedural steps of *SRSSD*, which comprises three stages:

S1 ***Matrix* H *update***. In the first stage, both **U** and **V** are fixed and **H** values are updated. As said above, one can update **H** by computing 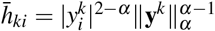. However, to avoid numerical instabilities near zero, we adopt the following smoothed update: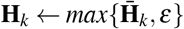 with *ε* « 1.

S2 ***Sparse Coding***. In the second stage, both **V** and **H** are fixed and **U** values are updated. Note that equation (3) comprises two terms to be minimized, but the second term does not depend on **U**. This implies that the optimization problem of equation (3) can be reformulated as: 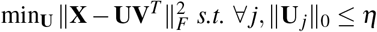. There are several well-known “pursuit algorithms” that find approximate solutions to this kind of problems, such as Basis Pursuit (BP) and Orthogonal Matching Pursuit (OMP)^61^. Here, we approximate the *ℓ*_0_ norm (a non convex problem) with its best convex approximation, i.e., *ℓ*_1_ norm – which allows us to perform sparse coding by applying an Iterative Soft-Thresholding (IST)^62^. Note that this stage is equivalent to the application of Algorithm 2, where we apply only sparse coding with a fixed dictionary.

S3 ***Structured Atoms***. In the third stage, both **U** and **H** are fixed and the dictionary **V** is updated; furthermore, following^35^, an atom structured sparse representation can be found. As both **U** and **H** are fixed, the problem of equation (3) can be reformulated as:

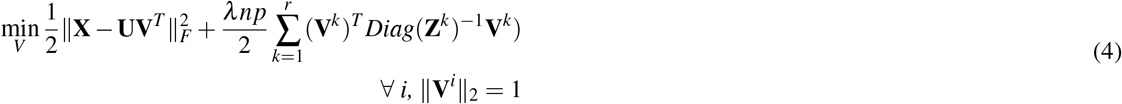

In this case, as both terms of Equation (4) are convex and differentiable terms with respect to **V**, a closed-form solution for **V** can be found. However, a proximal method is considered to avoid *p* matrix inversions:

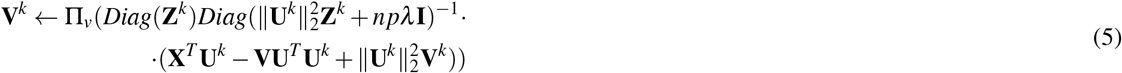

where Π_*v*_(*w*) is simply the Euclidean projection of *w* onto the unit ball, and the argument of Π_*v*_ is obtained by composing a forward gradient descent step on the first term with the proximity operator of the second term of (4).

### Method for reconstructing test trajectories and calculating reconstruction error (RMS)

To build the dataset to be reconstructed, we split the trajectories into training segments or patches *X* ∈ ℝ^*n*×*p*^, where *n* is the number of patches (10000 in our test set) and *p/*2 is the length in time of the patches. Each row of *X* includes a sequence of x and y coordinates that compose a patch, e.g., x1, y1, x2, y2, etc. We split trajectories lasting more than *p/*2 time steps into two or more rows of *X*. Then, we use SRSSD to learn a dictionary *V* ∈ ℝ^*p*×*r*^ of *r* atoms, having maximum length *p*. For the sake of comparison, we adopt the same procedure using PCA (where the matrix *V* represents *r* principal components; but note that PCA requires all element to be the same length, which we fix to *p*).

In general, once learned the Dictionary *V*, *X* can be reconstructed by computing *U* ∈ ℝ^*n*×*r*^, where *U* represents the coefficients of the atoms computed with sparse coding (see Algorithm 2), or the PCA coefficients *U* = *XV*. Using the coefficients, a new reconstruction matrix *X_rec_* = *UV^T^* (with *X_rec_* ∈ ℝ^*n*×*p*^) can be obtained, which can be compared with the original *X*, to calculate a reconstruction error RMS. However, this method would only measure how well the learned dictionary represents the learned trajectories, not how well it generalises.

For this, we use a more compelling methodology, inspired by the *missing pixel method*: we create “holes” in the original trajectories (by removing consecutive columns of *X*) and measure how well our method permits to reconstruct them. More specifically, we firstly build a *restricted* matrix 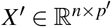, with *p*’ < *p* by removing *p* − *p*’ (consecutive) columns from *X*. Therefore, we build a *restricted* dictionary 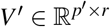 (o *r* PCA components), by removing *p* − *p*’ rows from *V*. We then compute the matrix *U*’ ∈ *R*^*n*×*r*^, using *X*’ and *V*’ (instead of *X* and *V* as above). Because *U*’ has the same matrix dimensions as *U*’, it is possible to use the original matrix *V* to compute 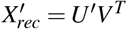, with 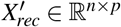 that has the same dimension of original matrix *X*, thus resulting in a “full” trajectory with reconstructed holes.

Finally, we calculate the reconstruction error RMS using a *Frobenius* norm ‖·‖_*F*_, which considers the difference between a target trajectory *X* ∈ ℝ^*n*×*p*^ and 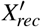.

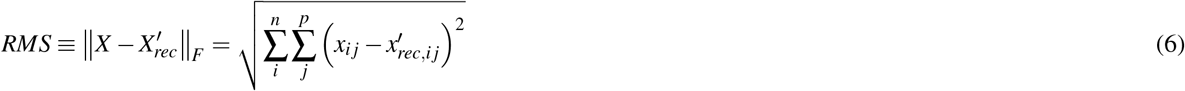

We adopt the same (missing pixel) procedure for both SRSSD and PCA, for test and validation. Furthermore, to test the robustness of the method, we use the learned dictionaries to reconstruct different portions of trajectories, from 10% to 90% – the latter coming closer to reconstructing full trajectories.

**Algorithm 1.**
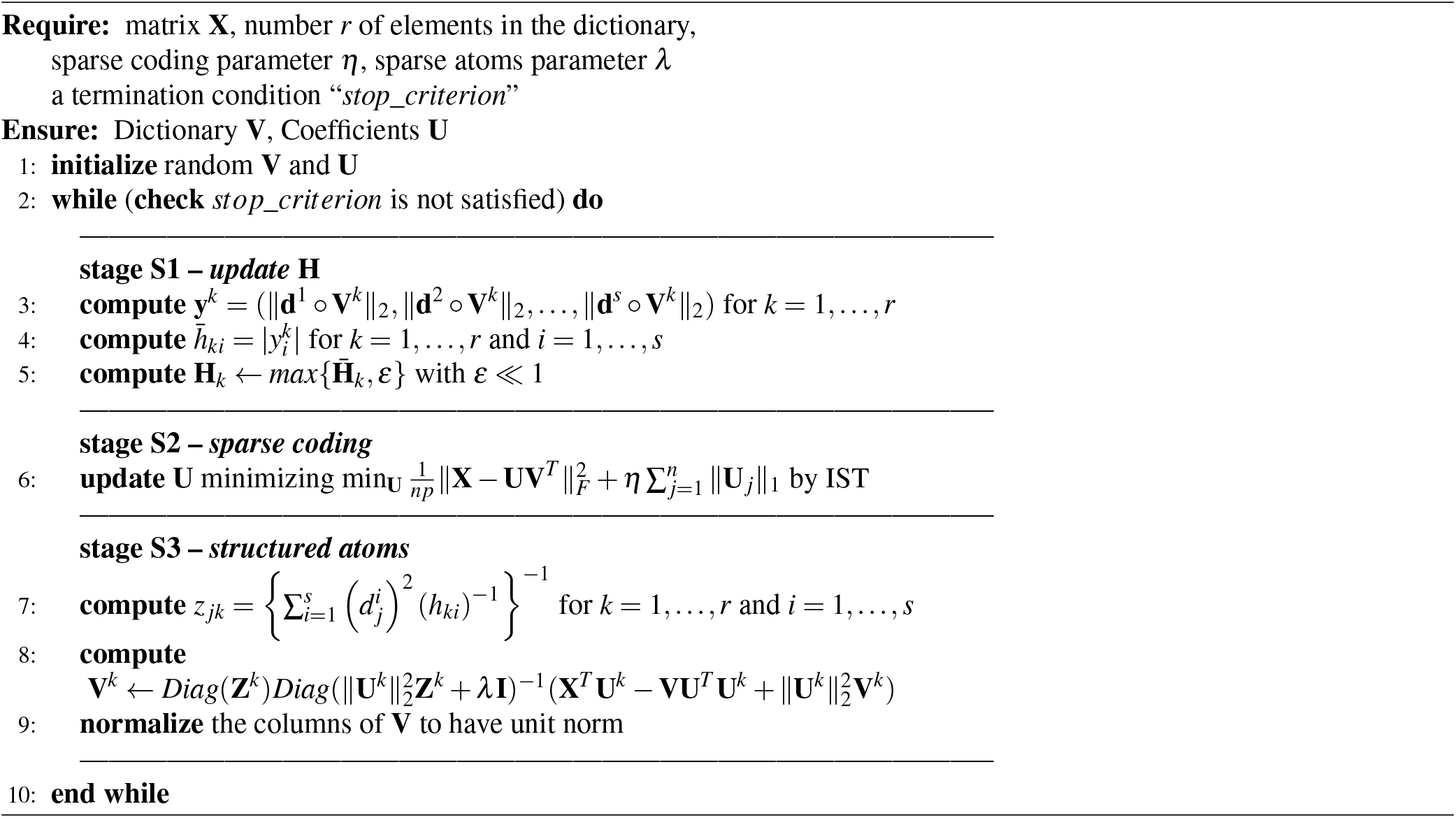
*SRSDD*(**X***, r, η, λ, stop*_*criterion*)

**Algorithm 2.**
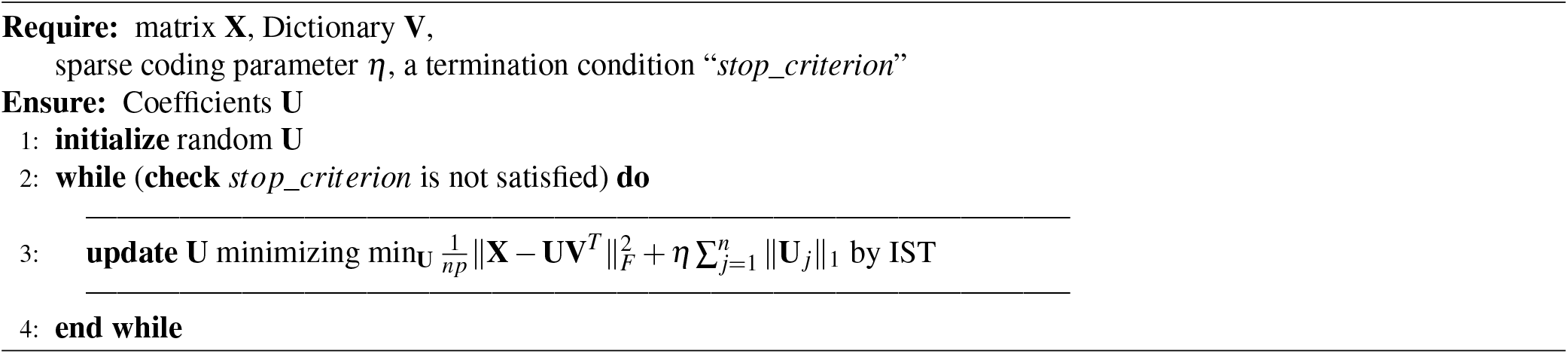
*Sparse Coding*(**X**, **V***, η, stop*_*criterion*)

## Acknowledgments

This research has received funding from the European Union’s Horizon 2020 Framework Programme for Research and Innovation under the Specific Grant Agreements No. 785907 and 945539 (Human Brain Project SGA2 and SGA3 to GP), the European Research Council under the Specific Grant Agreement No. 820213 (ThinkAhead to GP) and the Human Frontier Science Program (grant no. RGY0088/2014 to GP, MVDM and CK). The GEFORCE Titan GPU card used for this research was donated by the NVIDIA Corp. The funders had no role in study design, data collection and analysis, decision to publish, or preparation of the manuscript.

## Author contributions statement

F.D, R.P., M.v.d.M., C.K. and G.P. conceived the experiment(s), F.D., R.P., D.M., S.F., E.I. and G.P. conducted the experiment(s). All authors reviewed and analysed the manuscript.

## Notes

### Competing Interest Statement

The authors have declared no competing interest.

