## Supplemental information for "A framework to identify structured behavioral patterns within rodent spatial trajectories"

### Maze complexity measures

The figures shown below illustrate the results of maze discretization (binning), as used to measure maze spatial complexity (Figure S1); occupancy of animal R01 in the third phase of each day, using this binned space (Figure S2), the time required to complete a trial and secure a reward (Figure S3), and example trajectories during phase 3 of day 1 (Figure S4).

#### Experimental results, animals R02 and R03

In the main text, we have focused on animal R01 for illustrative purposes. Our results generalize to the two other two animals (R02 and R03) in our dataset. Figures S5 and S6 shown below illustrate that 1) reconstruction error is lower using SRSSD compared to PCA in both animals R02 and R03; 2) in both animals, coefficient sparsity correlates with measures of spatial complexity of the mazes (compare to Figure 7 in the main article).

Figure S7 shows that coefficient sparsity of animals R02 and R03 increases in phase 1 of day 1, while the animals are re-familiarizing themselves with the maze, as in R01 (discussed in the main text). Figures S9 and S9 show that classification of (U-path or shortcut) trajectories is accurate for R02 and R03, respectively, akin to R01 (discussed in the main text).

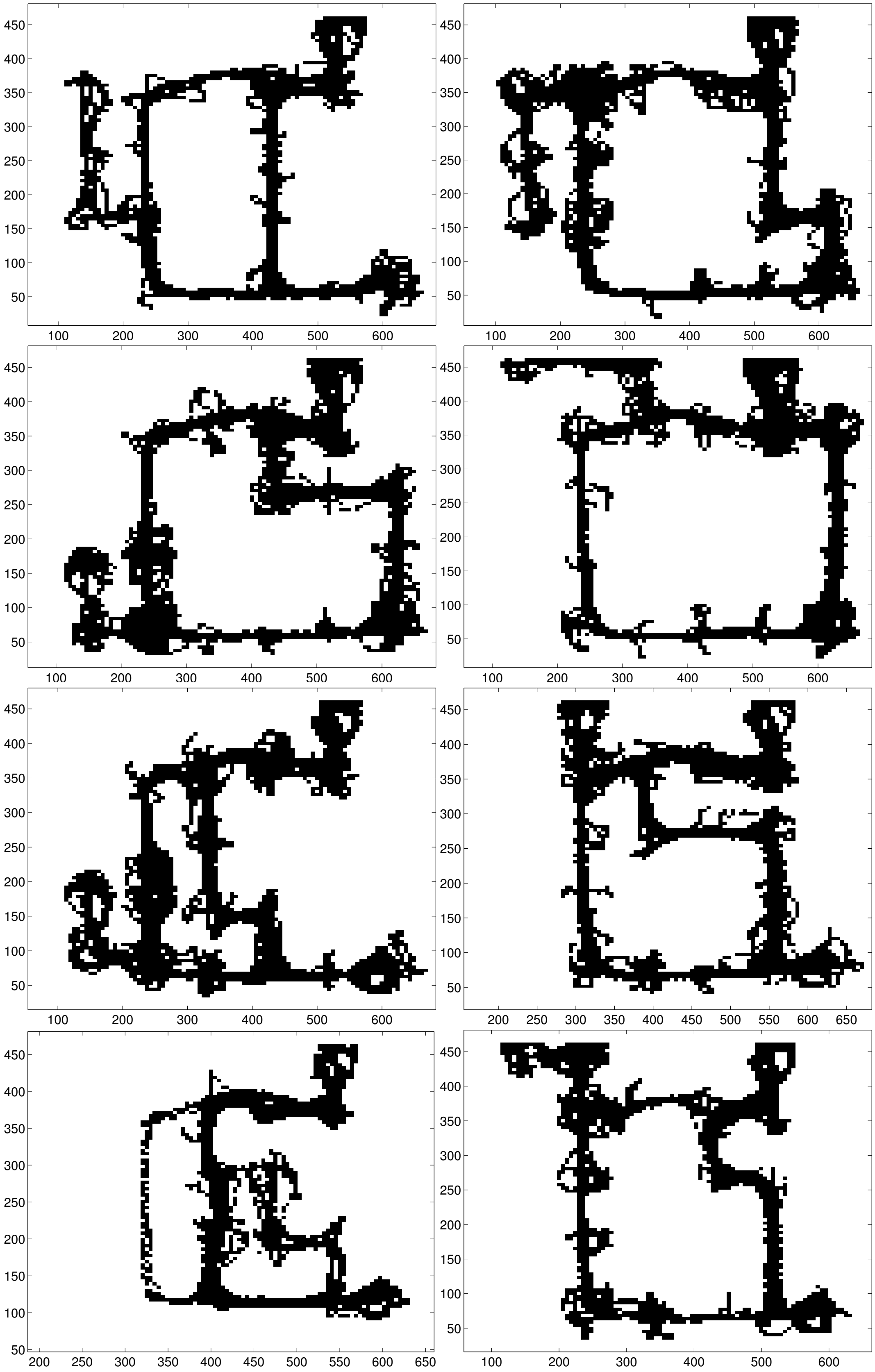


**Figure S1.** Discretization of mazes into bins used to measure maze spatial complexity

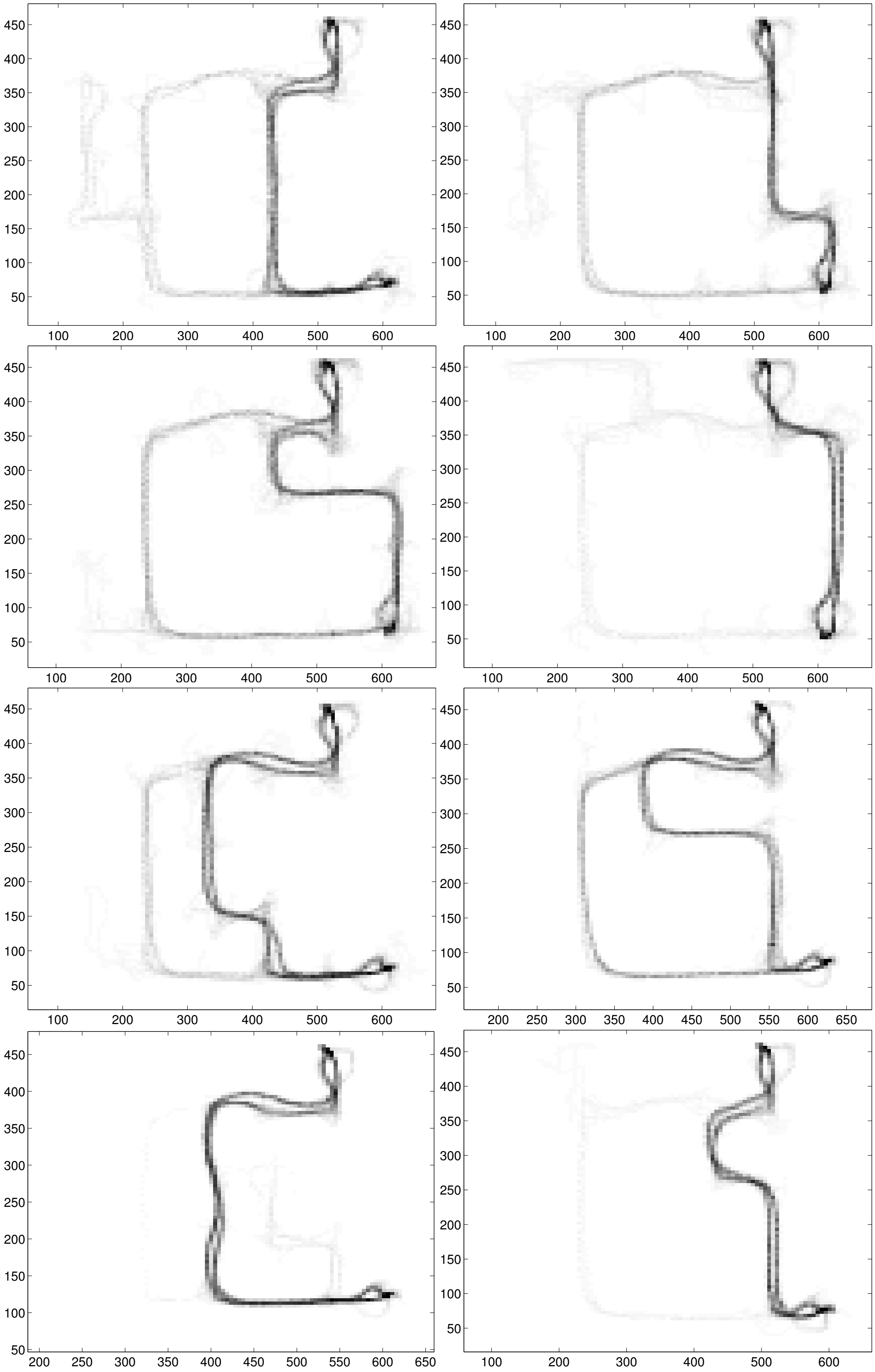

**Figure S2.** Occupancy of R01 in the third phase of each day, after binning the space.

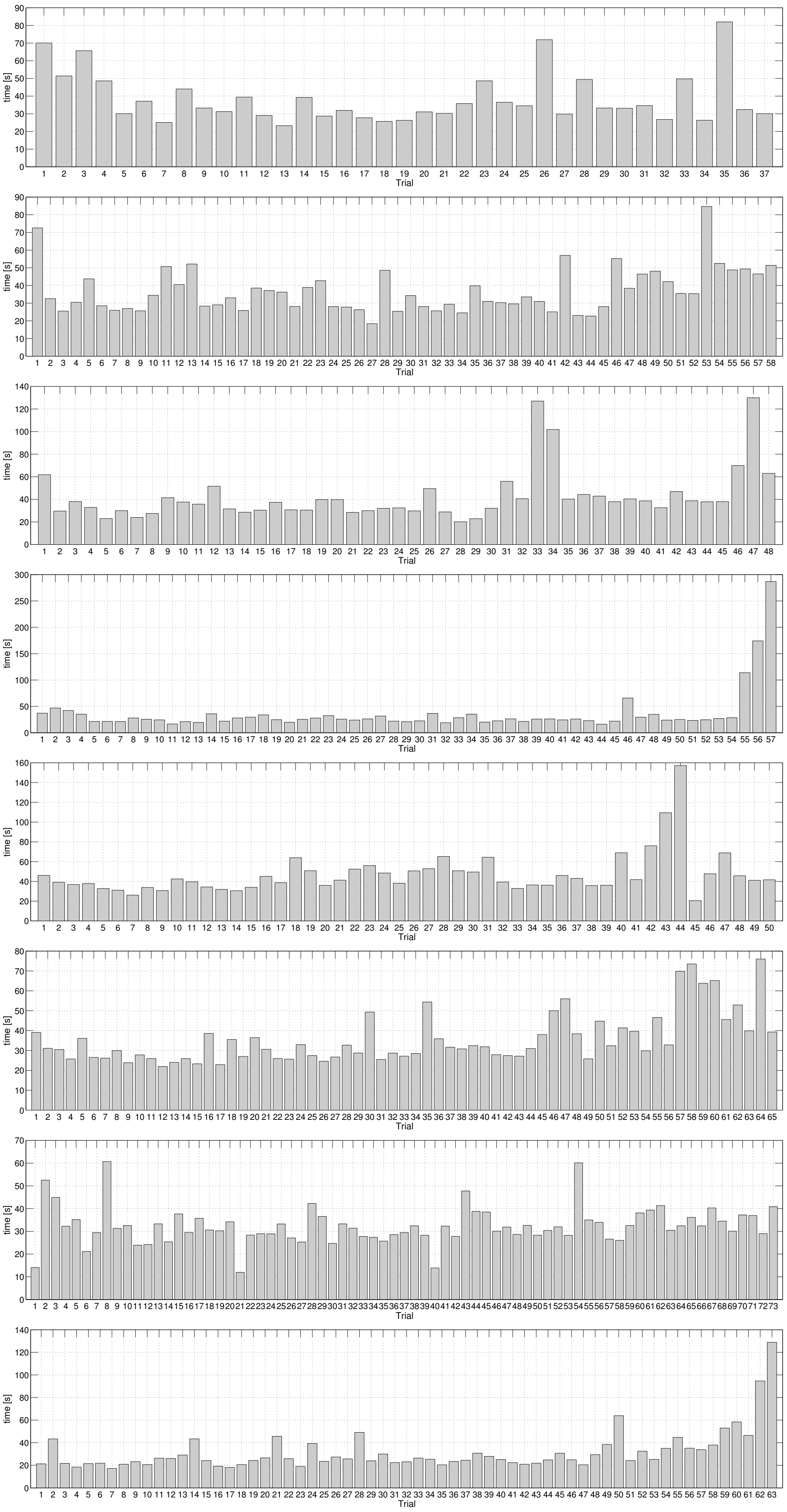


**Figure S3.** Time required for completing a trial (and collecting a reward) for each day, animal R01

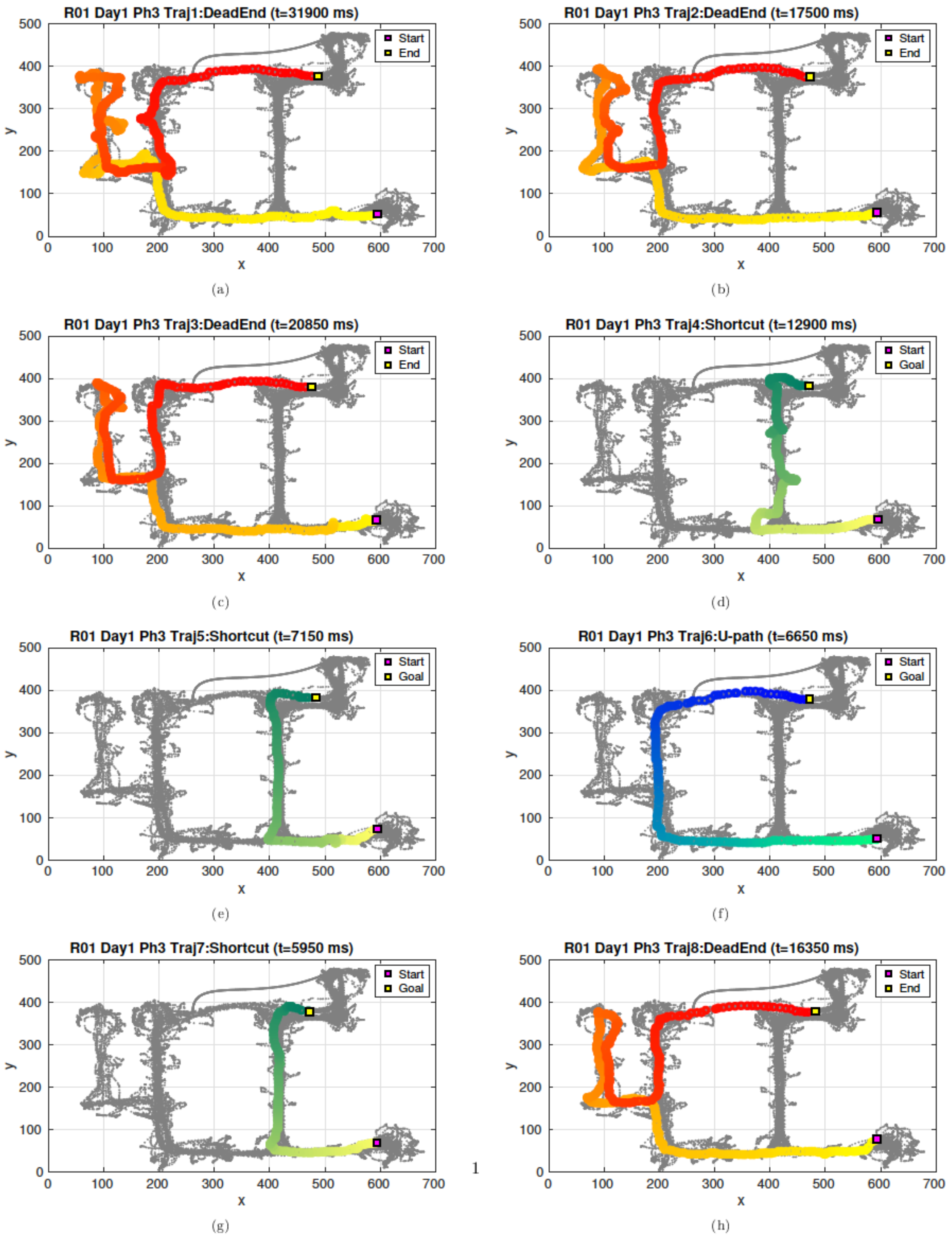


**Figure S4.** Eight examples of trajectories classified as U-path, shortcut or dead-end, during day 1, phase 3, in animal R01

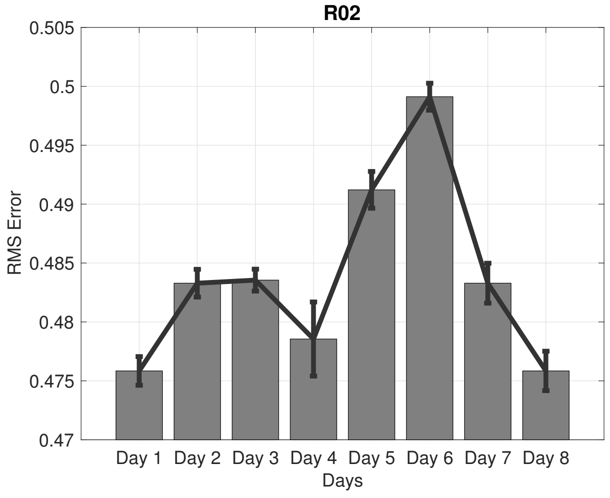


**(a)** RMS Error

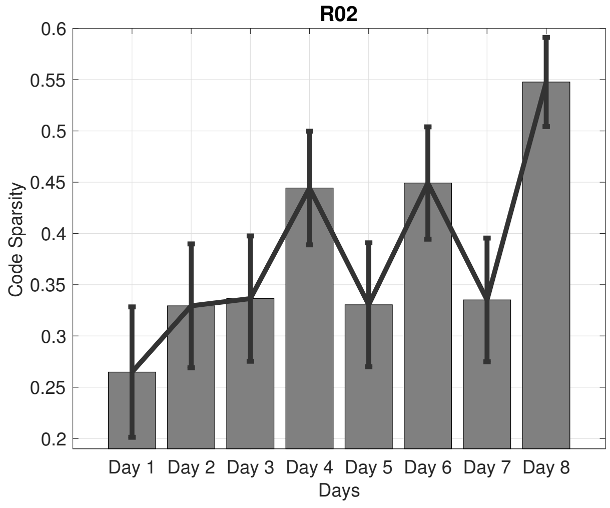


**(b)** Coefficient Sparsity

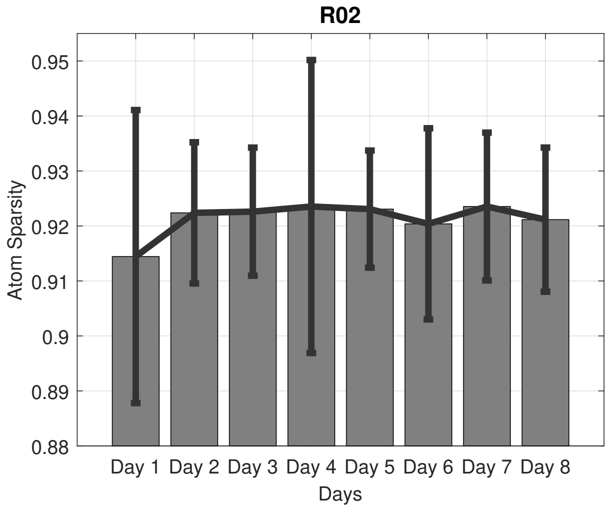


**(c)** Atom Sparsity

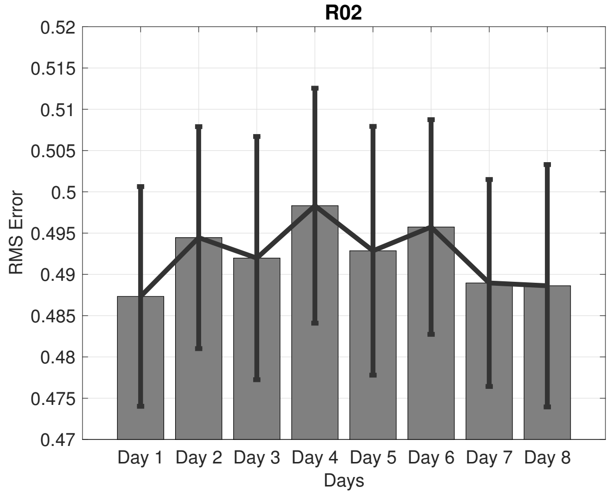


**(d)** Baseline

**Figure S5.** Reconstruction across 8 days of R02 (after learning only from data of day 1. Noise is 0.5 and the best 30 SRSSD dictionaries found on day 1 are used. (a): RMS Error. (b): coefficient sparsity. (c) dictionary sparsity. (d) Baseline performance: reconstruction of Phase 1.

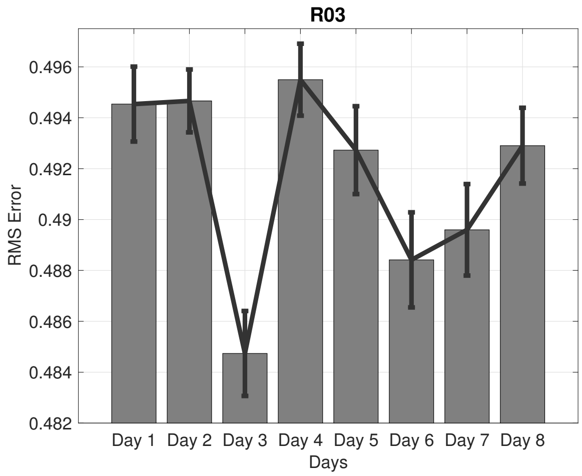


**(a)** RMS Error

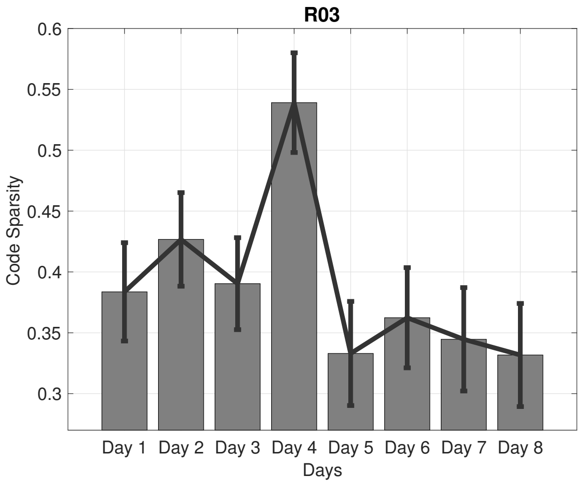


**(b)** Coefficient Sparsity

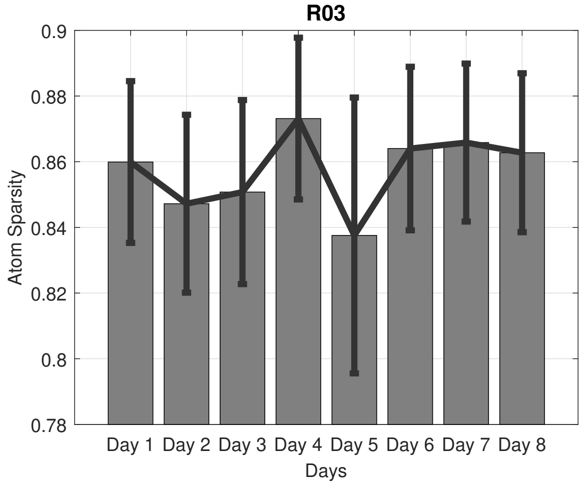


**(c)** Atom Sparsity

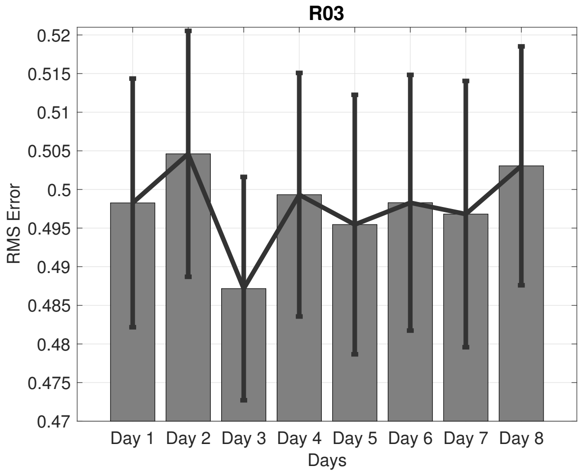


**(d)** Baseline

**Figure S6.** Reconstruction across 8 days of R03 (after learning only from data of day 1. Noise is 0.5 and the best 30 SRSSD dictionaries found on day 1 are used. (a): RMS Error. (b): coefficient sparsity. (c) dictionary sparsity. (d) Baseline performance: reconstruction of Phase 1.

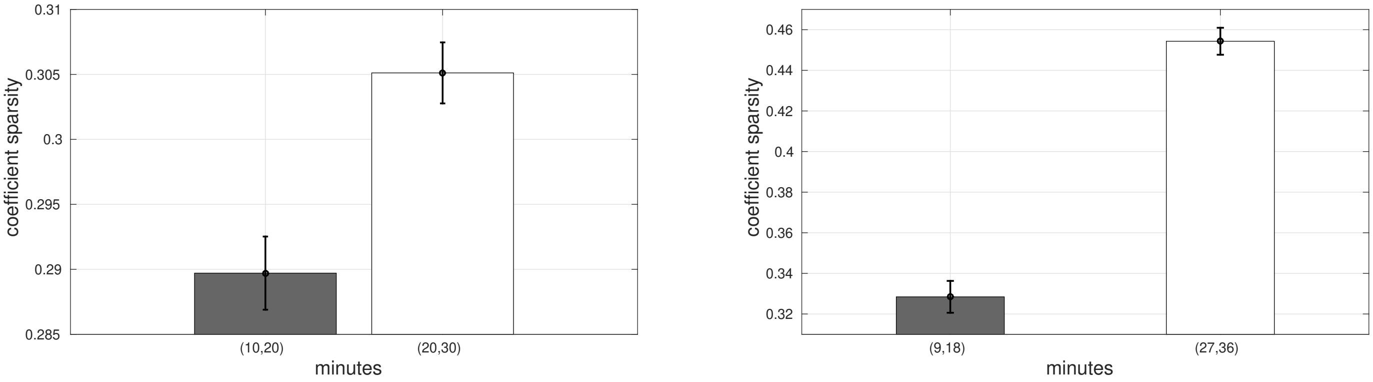


**(a)** Coefficient sparsity, R02 **(b)** Coefficient sparsity, R03

**Figure S7.** Coefficient sparsity of animals R02 and R03 increases in phase 1 of day 1, while the animals learn navigating the U maze, thus revealing stereotypy of behaviour. We report the comparison between the 2nd and 4th intervals of the same length.

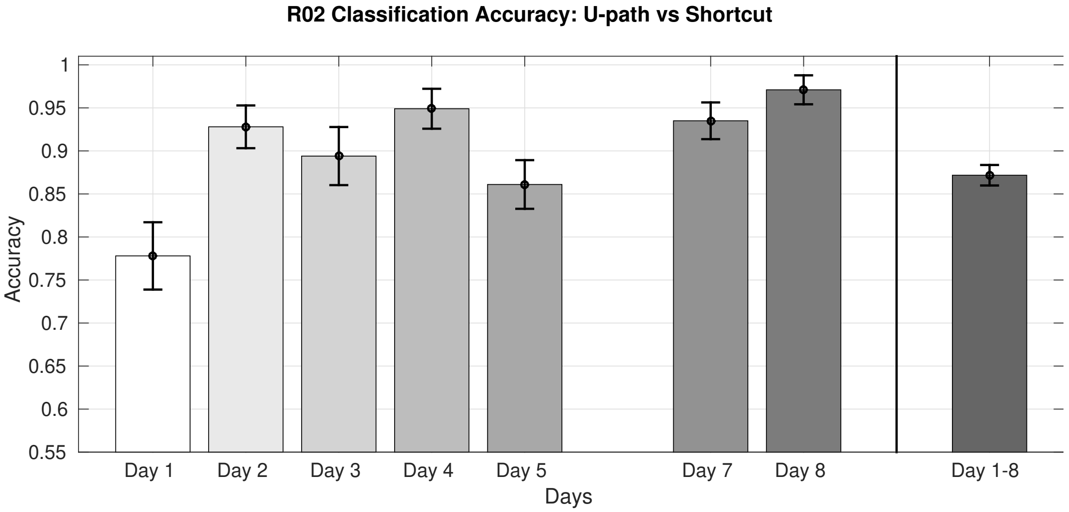


**Figure S8.** Classification of trajectories (U-path or shortcut) of phase 3, day 1, animal R02. The score for day 6 is missing, since on that R02 always selects the shortcut.

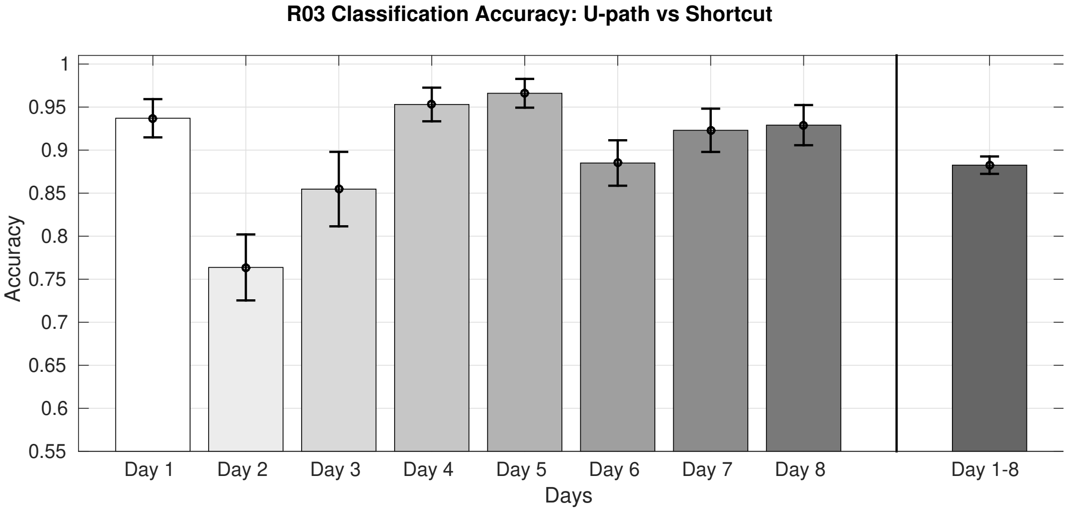


**Figure S9.** Classification of trajectories (U-path or shortcut) of phase 3, day 1, animal R03
